# Dynamical buffering of reconfiguration dynamics in intrinsically disordered proteins

**DOI:** 10.1101/2025.10.12.681911

**Authors:** Miloš T. Ivanović, Andrea Holla, Mark F. Nüesch, Valentin von Roten, Benjamin Schuler, Robert B. Best

## Abstract

The dynamics of intrinsically disordered proteins are important for their function, allowing their heterogeneous conformational ensembles to rapidly reconfigure in response to binding partners or changes in solution conditions. However, the relation between sequence composition and chain dynamics has rarely been studied. Here, we characterize the dynamics of a set of 16 naturally occurring disordered regions of identical chain length but with highly diverse sequences. In spite of the strong variation of chain dimensions with sequence in this set inferred from single-molecule FRET, nanosecond fluorescence correlation spectroscopy yields chain reconfiguration times that are almost independent of sequence. This surprising observation contrasts with the slowdown in dynamics, attributed to internal friction, that has been observed in more compact disordered proteins. We investigated this effect with the aid of multi-microsecond, all-atom explicit-solvent simulations of all 16 disordered proteins. The simulations reproduce the experimental FRET efficiencies with near-quantitative accuracy, with explicit inclusion of the FRET dyes improving agreement with experiment while minimally perturbing the protein ensemble. Critically, the simulations also reproduce the lack of correlation between reconfiguration times and chain dimensions across the sequences and allow us to rationalize this observation as arising from two competing factors as the chains get more compact. The narrowing of end-to-end distance distributions and a concomitant reduction of the corresponding intrachain diffusion coefficients have opposite effects that end up resulting in only a small variation of reconfiguration times with chain dimensions. These compensating factors “buffer” the effect of sequence on linker dynamics, which may help to conserve function as sequences evolve.

## Introduction

Intrinsically disordered proteins (IDPs) and intrinsically disordered regions (IDRs) are now recognized to play a broad range of functional roles in biology, including molecular signaling, transcription regulation, and formation of biomolecular condensates. ^1–3^ They are also implicated in aberrant behaviour, as most amyloids are formed by disordered proteins,^4,5^ and “aging” of the disordered regions within condensates has been associated with disease outcomes such as ALS.^6,7^ Disordered linker sequences have also been shown to play an important role in maintaining an appropriate distance between the folded domains which they bridge.^8^

A key feature of disordered regions enabling their function is the extremely broad distribution of configurations which they populate, compared to folded domains which can usually be modeled as a single structure with only local fluctuations – such flexibility underlies the ability of IDRs to bind alternate partners or to form disordered complexes, for example. ^9^ Equally important for function is the rapid interconversion between configurations in the disordered ensemble, often characterized in terms of a reconfiguration time.^10^ However, these broad ensembles and rapid dynamics present a challenge for studying IDRs and IDPs experimentally. Techniques which can provide information on structure and dynamics of disordered ensembles include NMR,^2^ small angle X-ray scattering (SAXS), ^11^ Förster resonance energy transfer (FRET),^10,12^ nanosecond fluorescence correlation spectroscopy (nsFCS)^10,13^ and neutron spin-echo experiments. ^14^ Even for these methods, however, the limited number of independent observables and signal averaging over a heterogeneous ensemble limit the detailed interpretation of these experiments.^15,16^ In this context, molecular simulations with either all-atom or coarse-grained force fields can play a critical role in filling in the structural and dynamic details.^17,18^ Without constraints from experiment, however, the results of such simulations are dependent on the accuracy of the underlying simulation force field.^17^ A close combination of the above experiments together with molecular simulations can therefore provide a powerful strategy for decoding the dynamic structural ensemble of IDRs. ^15,16^

Despite the importance of the reconfiguration dynamics of IDRs, relatively little has been done to study its sequence dependence systematically. In this work, we address this deficiency by investigating a set of intrinsically disordered linker regions from RNA-binding proteins, carefully selected to have diverse sequence properties and whose FRET efficiencies have previously been determined.^19^ We probe the corresponding dynamics of these 16 IDRs, by measuring their chain reconfiguration times, *τ*_*r*_, via nanosecond fluorescence correlation spectroscopy (nsFCS). This method allows us to measure the end-to-end distance dynamics by means of the concomitant fluorescence fluctuations of the donor and acceptor dyes.^10,13^ We observe very rapid chain dynamics, with values of *τ*_*r*_ scattered around ∼ 30 ns, in the range expected for IDPs of this chain length.^10^ Remarkably, despite the broad range of chain dimensions of these IDRs indicated by their FRET efficiencies, their reconfiguration times are remarkably similar for all sequences and surprisingly uncorrelated with their FRET efficiency – in contrast to the slowdown of dynamics that might be expected from internal friction effects with increasing compaction. ^9,20–22^ To complement these results and investigate the relation between chain dimensions and dynamics in mechanistic detail, we have performed multi-microsecond all-atom explicit-solvent simulations of the entire set of linker proteins, both with and without the FRET chromophores explicitly represented. The simulations reproduce the experimental trends in FRET efficiency almost quantitatively, and the chromophores only minimally perturb the conformational ensemble; however, inclusion of the chromophores does yield FRET efficiencies in closer accord with experiment. We also find that the narrow range of reconfiguration times and lack of correlation with average FRET efficiencies seen in experiment is reproduced by the simulations. We then use the equilibrium and dynamic properties accessible in the simulations to rationalize the lack of correlation between reconfiguration times and FRET efficiencies by approximating the dynamics of the interchromophore distance as one-dimensional diffusion. As we show, compensating effects arising from the sequence-dependent variation of the end-to-end diffusion coefficient on the one hand and of chain compactness on the other lead to the approximately sequence-independent reconfiguration times. We further cross-validate our explanation against the available experimental data and discuss the implications for IDR function.

## Results

### nsFCS experiments yield very similar reconfiguration times for a diverse set of linker IDRs

We investigate the effect of IDR sequence on reconfiguration time using a sequence-diverse set of intrinsically disordered linker regions from RNA-binding proteins^19^ (Table S1). The previously determined FRET efficiencies for these linker IDRs range from roughly 0.4 to 0.9 for the dye pair Cy3B/CF660R (Fig. 1A,B), indicating that these sequences indeed encode a broad range of chain expansion. ^19^ Their polymer scaling exponents *ν* inferred from the SAW-*ν* model^23^ range from values slightly above 3/5, close to the expectation for a self-avoiding walk (no attractive interactions), to slightly below 1/2, the value expected when attractive and repulsive interactions are balanced. However, none of the chains approach the compaction of collapsed globules with *ν* ≈ 1*/*3. Examples of distance distributions and configurations corresponding to the most expanded and most collapsed variants are shown in Fig. 1B, illustrating the substantial range of compaction sampled.

**Figure 1.**
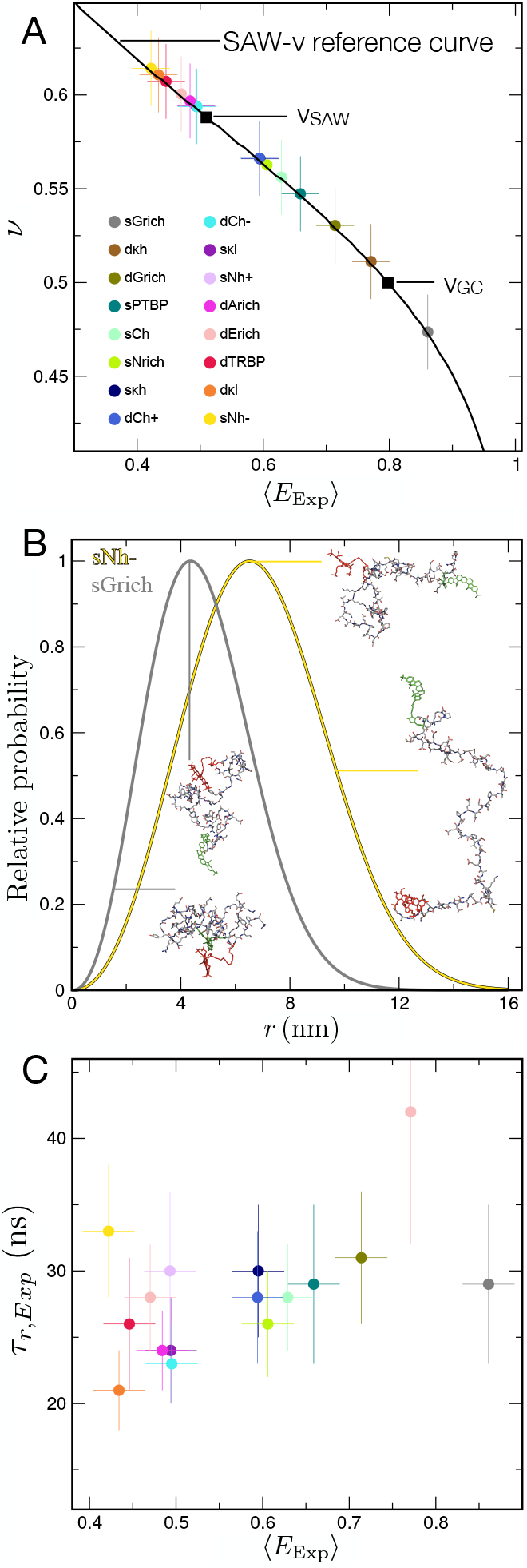
Chain dimensions in a set of linker IDRs with diverse sequence properties. (A) Scaling exponents, ν, inferred from experimental FRET efficiencies, ⟨*E*_*Exp*_ ⟩, using the SAW-*ν* model ^23^ show linkers ranging from close to self-avoiding walk (SAW) chains to Gaussian chain (GC)-like. The uncertainty in the FRET efficiencies is taken as 0.03, and the uncertainty in the scaling exponents as 0.02. ^19^ The naming of the different sequences (see legend) reflects the sequence properties (see Table S1 and Holla et al. ^19^). (B) Examples of SAW-*ν* end-to-end distance distributions *P*(*r*) inferred from the experimental FRET efficiencies, contrasting the most expanded (sNh −) and most collapsed (sGrich) chains, with illustrative configurations from the simulations. Hydrogen atoms are omitted for clarity. (C) Lack of correlation of experimentally measured chain reconfiguration times, τ_*r,Exp*_, with the observed FRET efficiencies. The uncertainty of the chain reconfiguration times is reported in the Table S2.

To complement the previously measured transfer efficiencies of the linker IDRs,^19^ we have probed the chain dynamics with nanosecond fluorescence correlation spectroscopy (nsFCS), which allows us to measure the chain reconfiguration time within the disordered ensemble.^10,13^ Briefly, the distance fluctuations between donor and acceptor dye cause fluorescence intensity fluctuations, which can be recorded with nsFCS and used to obtain the end-to-end distance correlation times of the linker IDRs (see Methods). The chain dynamics are exceedingly rapid, with reconfiguration times scattered around ∼ 30 ns. Based on previous work on the dynamics of disordered proteins, we had expected a variation of these chain reconfiguration times with sequence, because the sequences under consideration here span a wide range of *R*_g_, and internal friction — dissipative forces within proteins that slow down their dynamics^20,24,25^ — and tend to increase with chain compaction, slowing down chain dynamics.^10,20,22,26,27^ However, in contrast to these expectations, the correlation times show at most a weak correlation with the FRET efficiencies (correlation coefficient, obtained from Monte Carlo parametric bootstrapping, *ρ*_MC_ = 0.35 (0.21)) (Fig. 1C) and thus chain compaction.

### All-atom molecular simulations reproduce equilibrium FRET and SAXS data for the linker IDRs

To investigate the unexpected lack of correlation between mean FRET efficiency and reconfiguration time in more depth, we performed all-atom, explicit-solvent simulations of the entire set of linker IDRs described previously^19^ for over six microseconds each in the Amber ff99sbws force field^29^ and conditions matched to experiment. To compute FRET efficiencies from the simulations with the fewest possible assumptions, we have explicitly included the chromophores in the molecular model (for sequences and dye positions, see Table S1). Parameters for the dyes Cy3B and CF660R were derived for this work using standard procedures as described in Methods. A key adjustment for the chromophores was a scaling of the interactions between the chromophores and water, analogous to that used for the protein force field, as discussed previously.^30,31^

The mean FRET efficiencies calculated based on the distance between the chromophores are compared with the corresponding experimental efficiencies in Fig. 2A, showing them to be in good accord. The difference between simulation and experiment is within the calculated standard errors for eight of the sixteen proteins, close to the 68 % expected for Gaussian-distributed errors, suggesting that the simulations accurately capture the dependence of chain dimensions on sequence.

**Figure 2.**
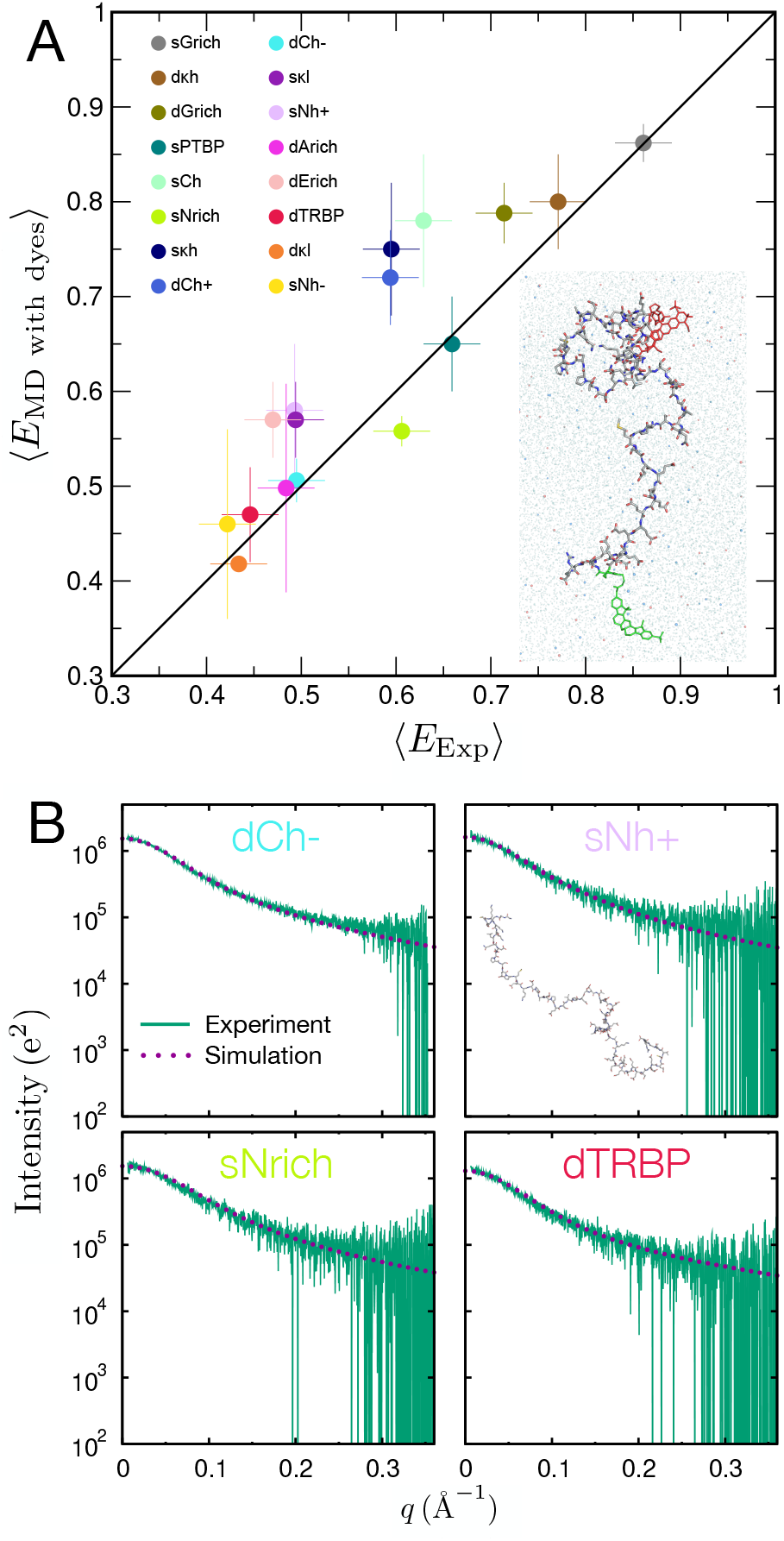
(A): Comparison between FRET efficiencies calculated from simulations with explicit chromophores and experimentally observed values (solid line: identity). Linear correlation coefficient and concordance correlation coefficient ^28^ are 0.90 and 0.84, respectively. Color code for IDRs is shown in the legend. The error bars represent one standard deviation estimated from the three independent simulation runs. Inset: A simulation snapshot showing the protein with explicit dyes, as well as the surrounding water and ions (only part of the simulation box is shown). The FRET donor and acceptor dyes are shown in green and red, respectively. K^+^ ions are shown as blue spheres and Cl^*−*^ ions as red spheres. Water molecules are shown as light blue, transparent spheres. Hydrogen atoms were omitted for clarity. (B): Comparison of the experimental SAXS curves with those calculated from the simulations (both without dyes attached).

As a further test of the simulations, we computed SAXS profiles for the four linker IDRs for which experimental data are available ^19^ (Fig. 2B, Fig. S1). Besides being a complementary technique to FRET, SAXS experiments can be performed without the need for conjugating the proteins to chromophores (we note, however, that SAXS experiments failed for several of the sequences owing to aggregation at the required concentrations ^19^). We also observe good agreement between the SAXS curves from experiment and from simulations, independently confirming the quality of the simulations.

Differences between the FRET efficiencies from experiment and simulation are close to expectations given the estimated errors. Nonetheless, we can assess potential sources of residual discrepancies between simulation and experiment from the relation between the deviation from experiment and various properties of the sequences under consideration, as plotted in Fig. 3A and Fig. S2. The strongest correlations with the deviation from experiment are to the fraction of charged residues (FCR) and the number of possible salt bridges, suggesting that the salt bridges in the simulations may be somewhat too strong, which has been highlighted for several force fields.^34–39^ We further tested this hypothesis by reweighting the simulations based on the total number of salt bridges, *N*_sb_, by assigning simulation frame *i* weight *w*_*i*_ ∝ exp[−*βϵ*_sb_], where *β* = 1*/k*_*B*_*T, k*_B_ is the Boltzmann constant, *T* is the temperature, and *ϵ*_sb_ is a test perturbation to salt bridge strength. The optimal perturbation of *ϵ*_sb_ ∼ 0.75 *k*_B_*T* (Fig. 3B) suggests that salt bridges in the current force field are slightly too strong. Using instead a correction with separate parameters for each type of salt bridge (Glu-Lys, Glu-Arg, Asp-Lys, Asp-Arg), *w*_*i*_ ∝ exp[−*β*(*ϵ*_EK_ + *ϵ*_ER_ + *ϵ*_DK_ + *ϵ*_DR_)], further improves the agreement with experiment yields *ϵ*_EK_ ≈ 1.29 *k*_B_*T, ϵ*_ER_ ≈ 3.75*k*_*B*_*T, ϵ*_DK_ ≈ 0.04 *k*_B_*T* and *ϵ*_DR_ ≈ 0.71 *k*_B_*T*, suggesting that any force field correction for salt bridges would likely need to account for the identity of the residues involved. The relatively small magnitude of the corrections required is consistent with the small differences from experiment in this work, and with the generally good agreement between experiments and simulations of complex coacervates of protein polyelectrolytes performed using the same force field. ^40,41^ The larger corrections involving arginine salt bridges may also help to explain why timescales for coacervates involving arginine-rich sequences are somewhat larger than the experimental values in simulations with this force field.^41^

**Figure 3.**
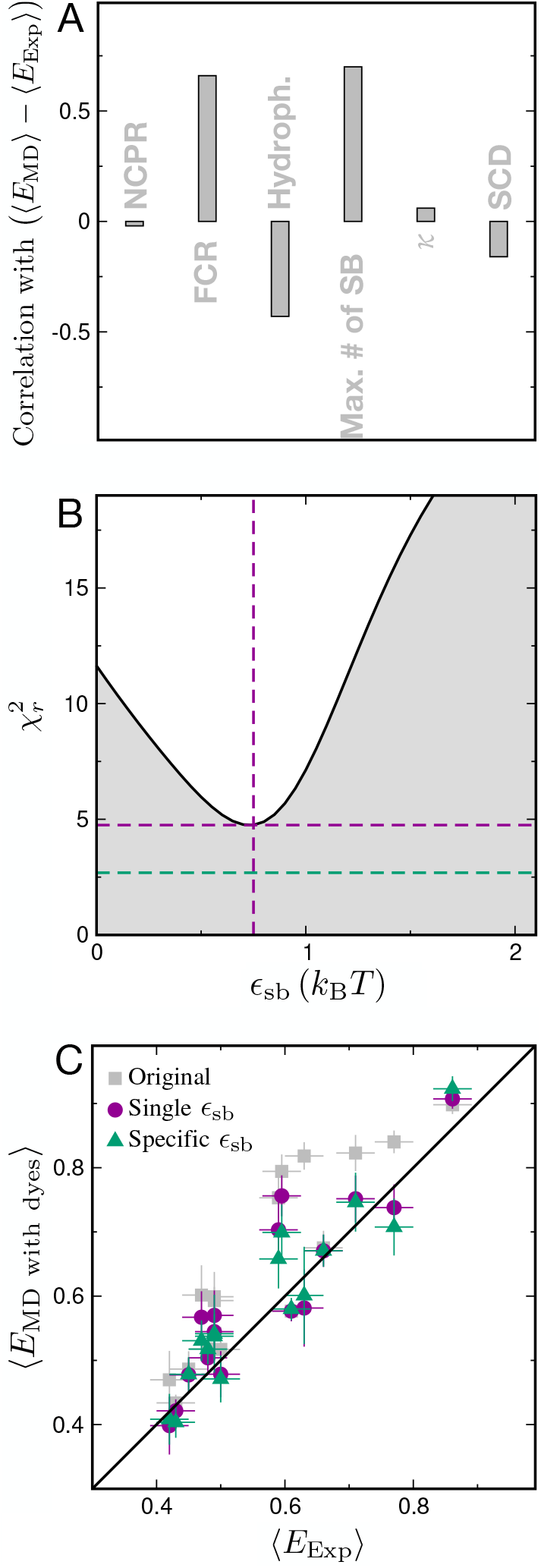
Reweighting of salt bridge interactions. (A): The correlation between the deviation of simulated and experimental FRET efficiencies and sequence properties commonly used to characterize disordered proteins: net charge per residue (NCPR), fraction of charged residues (FCR), hydrophobicity, number of possible salt bridges, *κ* parameter ^32^ and sequence charge decoration (SCD). ^33^ (B) and (C): The result of the optimization of a uniform salt bridge energy correction parameter, *ϵ*_sb_, used for reweighting the simulations based on the number of salt bridges. Effect of salt bridge energy corrections using a uniform parameter for all salt-bridges is shown in purple, and a correction using four individual parameters characterizing each possible type of inter-residue salt bridge is shown in green (see text for details).

It is worth noting that the residual discrepancies between simulations and experiments noted above for the more polyampholytic sequences may reflect not only slightly over-stabilized intramolecular salt bridges, but also remaining inaccuracies in the description of interactions between charged residues and their counterions.^42^

As an alternative reweighting approach we have also used a Bayesian method^43,44^ which has successfully been employed in a number of recent studies, ^45–47^ whereby each frame of the simulation is independently reweighted so that their average matches the experimental observables while also subject to a penalty term keeping the weights as close to uniformity as possible, employing as experimental data the mean FRET efficiency and the variance of the FRET efficiency distribution obtained from fluorescence lifetime information.^46^ As shown in Fig. S3 and Table S3, the reweighting is able to reproduce these data for each sequence to be within experimental error while still keeping the weights relatively uniform across the trajectory, a testament to the quality of the force field. The resulting equilibrium distributions and mean values of the end-end distance are thus only modestly perturbed from the original simulations; the most notable changes being to linkers sCh, s*κ*h and dCh+ (Fig. S3), as expected from the mean FRET efficiencies plotted in Fig. 2. Note that, for all 16 IDRs, the orientational factor *κ*^2^ is very close to the value of 2*/*3 (as expected for rapid isotropic rotational averaging of the dye transition dipoles, and in agreement with the low experimentally observed fluorescence anisotropies^19^) remains unaffected by reweighting (Table S3). We report the resulting weights together with the simulation ensembles in the repository associated with this work as a resource that we expect to be useful for the refinement of atomistic force fields or coarse-grained models.

### FRET chromophores minimally affect the dimensions of these IDRs

A common concern in FRET experiments is the potential perturbation caused by the extrinsic chromophores introduced for the purpose of measuring distance distributions. Clearly, addition of a probe must have some effect on the conformational distribution, and for some very hydrophobic probes, this effect has been clearly observed in experiment, ^12^ while in other cases the effect is more subtle.^19^ In our previous work on these linker IDRs, we had used a coarse-grained force field trained against experimental data that can empirically account for the small perturbations of conformational distributions caused by commonly used chromophores, which indicated that Cy3B/CF660R, the pair of chromophores we are using here, has weaker interactions with the protein than an alternate pair of dyes, the AlexaFluors 488 and 594.^19^ However, this force field optimization was based on a simplified “top-down” model driven only by experimental data, rather than by directly accounting for the detailed interactions involving the chromophores. To address the effect of the chromophores on the results, we thus performed all-atom simulations of all the linker IDRs, both with and without chromophores for a direct comparison. We estimated transfer efficiencies for the unlabelled proteins using the distance between the *α* carbon atoms of the labelled residues and scaling it to account for the effective increase in chain length from the cysteine side chains and dye linkers.^48^

We find that the inferred FRET efficiencies are very similar with and without the chromophores (Fig. 4A, Fig. S4), i.e., the effect of the dyes appears to be modest. However, there is an apparent shift to slightly lower transfer efficiency — i.e., larger distance — in simulations with explicit chromophores. This is certainly in contrast to suggestions that chromophores cause proteins to collapse.^49,50^ Note also that inclusion of the chromophores actually results in an improvement of the agreement with experiment (Fig. S4). To more directly compare the equilibrium ensembles with and without dyes, we computed the protein radii of gyration, *R*_*g*_, in both cases (excluding chromophores from the calculation, Fig. 4B), and find no difference between them outside of statistical error. One can therefore infer that the slight shift in dye-dye separation to longer distances relative to the end-end distribution of the unlabelled protein primarily reflects the anisotropic distribution of configurations adopted by the chromophores in the context of the protein (see Fig. S5), as expected from their excluded volume, rather than a perturbation of the protein ensemble by attractive interactions.

**Figure 4.**
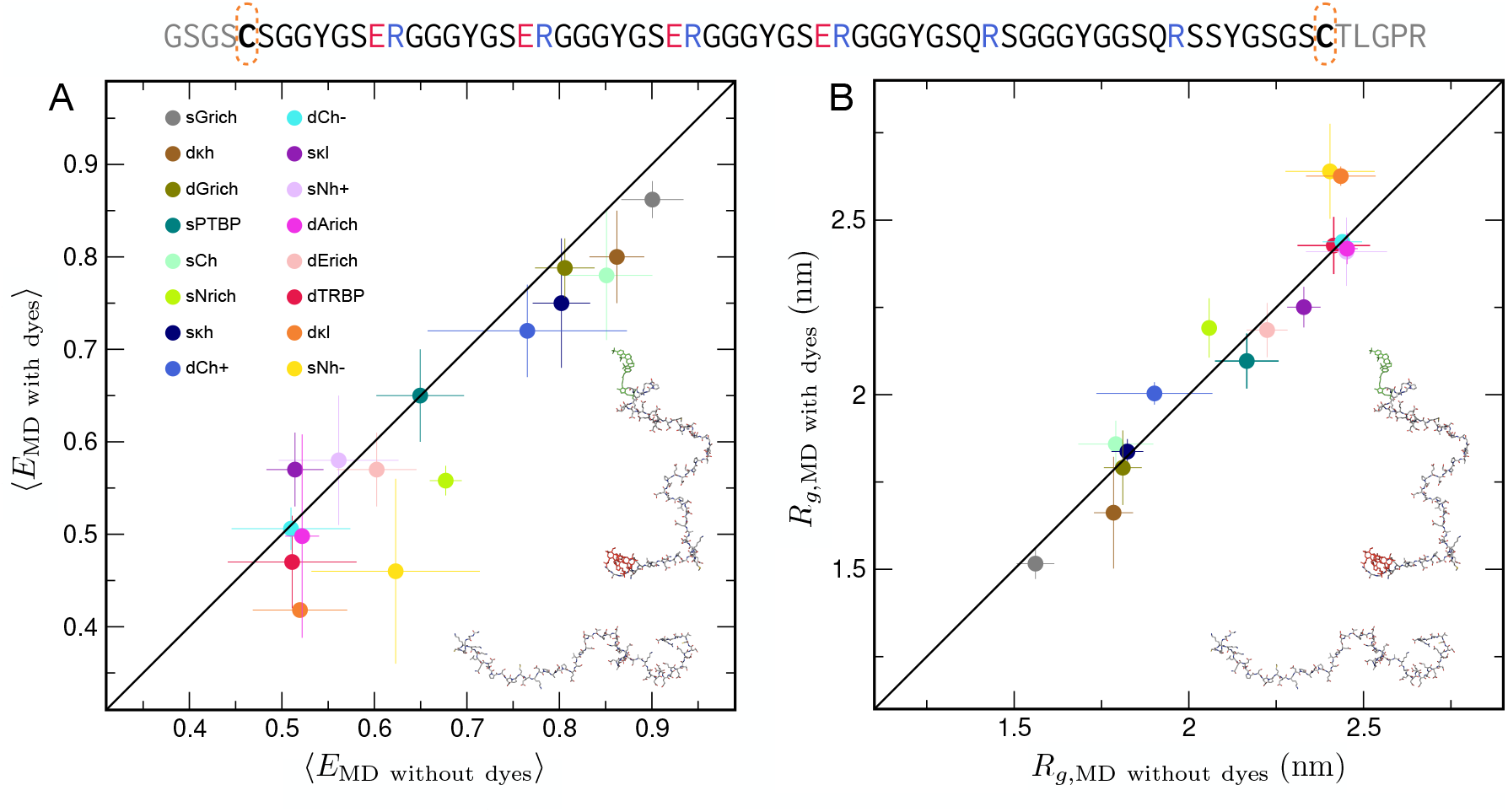
Effect of FRET dyes on protein configurations. (A) Comparison of average FRET efficiencies computed using the distances between the chromophore centers and the orientations between the dyes in simulations with FRET chromophores and the efficiencies estimated from simulations without dyes. ^48^ (B) Comparison of average protein radii of gyration (*R*_g_) from simulations with and without chromophores (chromophores excluded from *R*_g_ calculation). Error bars represent the standard deviations estimated from the three independent simulation runs. An example of one of the IDR sequences (sGrich) with the Cys residues used for labelling indicated is given at the top (see Table S1 for all sequences), and examples of snapshots of a dye-labeled and unlabeled linker IDR are shown as insets.

### Simulations explain lack of correlation between reconfiguration times and transfer efficiencies

Having shown that the simulations are consistent with the available experimental data reflecting the equilibrium configurational distribution, we turn to the central aspect, namely the chain dynamics, which can have a direct effect on binding kinetics and mechanisms.^51–53^

We analyze and interpret the simulations by using a one-dimensional diffusion model to describe the dynamics of the inter-chromophore distance, and we use the dynamical data from the three separate simulation runs of each linker to optimize the parameters of the model to match the distance distributions and chain dynamics observed in the trajectories.^54^ The optimal equilibrium distance distribution *P*(*r*) and distance-dependent diffusion coefficients determined from the model are shown in Fig. 5B and 5C, respectively. We have used the diffusion model to compute inter-dye distance correlation times for each protein, finding the values to lie in a similarly narrow range as that seen in experiment, and also that there is little correlation between reconfiguration times and mean FRET efficiencies, as in experiment. (Fig. 5A). In Fig. S6 we confirm that distance autocorrelation functions computed from this model are consistent with those estimated directly from the trajectories, with the advantage that the model correlation functions avoid the statistical noise seen in the long-time tails of those directly from the trajectories. As expected, the end-to-end distance distributions *P* (*r*) shift to larger distances for proteins with lower FRET efficiencies. Less obviously, the diffusion coefficients tend to decrease at shorter distances, a phenomenon which is common to all the linker proteins, and which has been observed earlier in simulations of model peptides: ^55^ this may be considered one manifestation of internal friction effects. ^10,27^ Although dissecting the individual contributions to internal friction is complex, in earlier work we have identified a slowdown in local reconfiguration dynamics via constraints on the chain and an increase in intrachain interactions as the primary sources of internal friction in collapsed chains. ^22,27^

**Figure 5.**
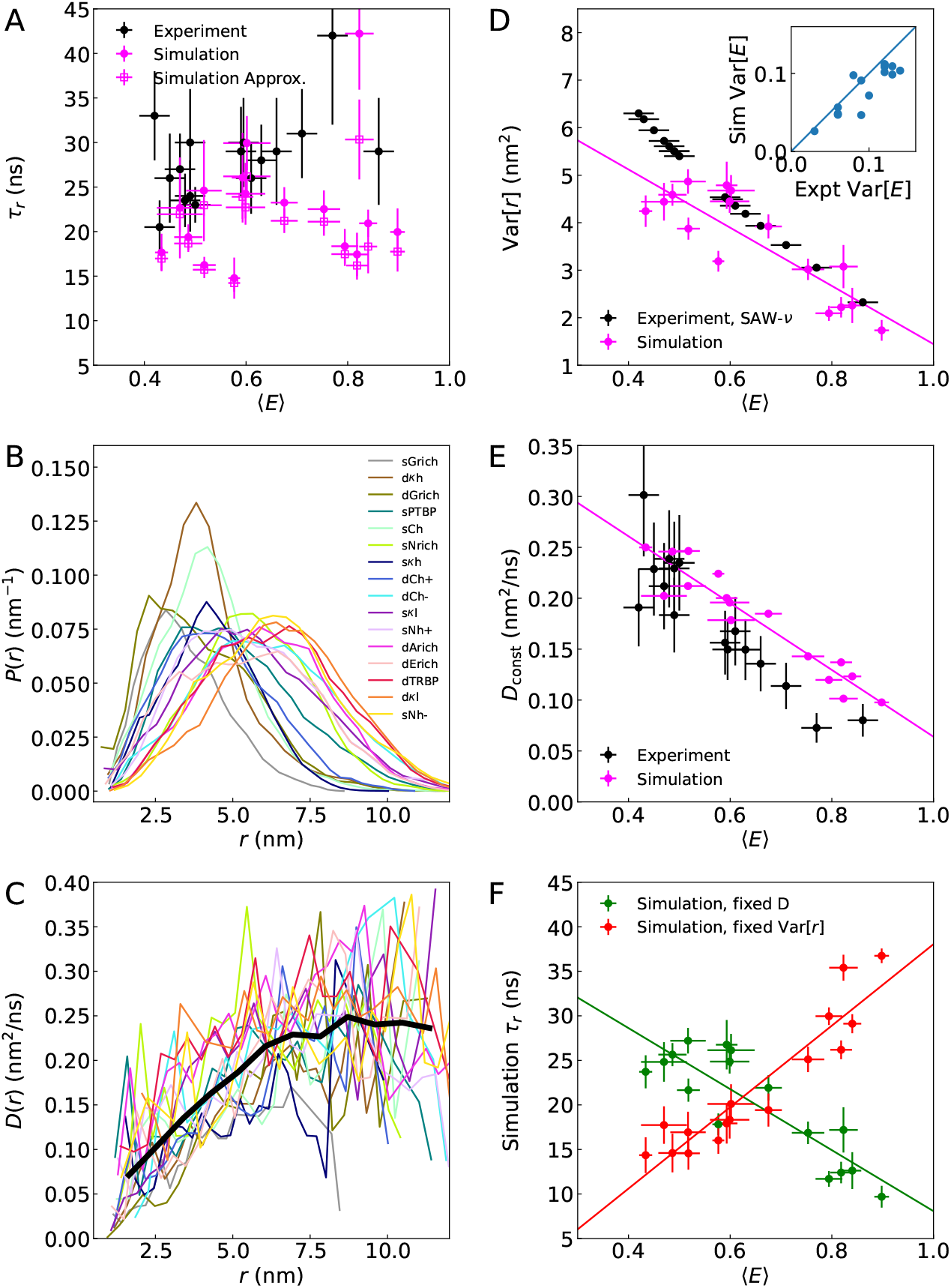
Rationalizing the lack of correlation between chain reconfiguration times and FRET efficiencies. (A) Weak correlation between reconfiguration times derived from nsFCS and FRET efficiencies from experiment (black, ρ_MC_ = 0.35 (0.21)), and between donor-acceptor distance autocorrelation times and FRET efficiencies in simulation (pink, ρ_MC_ = 0.19 (0.12)). Distributions of end-end distances (B) and position-dependent diffusion coefficients (C) inferred from a one-dimensional diffusion model applied to the simulation data. The thick black line in (C) represents the position-dependent diffusion coefficient averaged over all protein sequences. (D) Correlation between variances of end-to-end distance distributions and transfer efficiencies from simulation (pink), and between variances of distances inferred from SAW-*ν* model and transfer efficiencies from experiment (black). (inset) Correlation between Var[*E*] obtained from simulation with that from experimental lifetime measurements. (E) Correlation between position-independent diffusion coefficients and transfer efficiencies from simulation (pink) or experiment (black). Diffusion coefficients from simulation are inferred via optimization of a 1D diffusion model, while from experiment, *D*_const_ = Var_SAW−ν_ [*r*]*/*τ_*r*_. (F) Correlation times estimated from simulation via τ_*c*_ = Var[*r*]*/D*_const_, showing strong correlations if either Var[*r*] or *D*_const_ is fixed to the average value over all proteins from the simulations (ρ_MC_ = 0.89 (0.02) and −0.82 (0.04), respectively).

Position-dependent diffusion coefficients decreasing with *r* would be expected to lead to slower dynamics for more compact chains, as each protein is most sensitive to the value of *D*(*r*) at the most populated regions of *r*, given by the shape of *P* (*r*). Why is this not the case? A possible explanation can be simply described using the expression for the correlation time *τ*_*r*_ in a one-dimensional harmonic potential with constant diffusion coefficient *D*_const_,

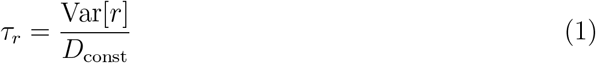

where Var[*r*] is the variance of the distance coordinate *r*.^54^ One can immediately see that changes both in the diffusion coefficient and in the variance, or width, of the distribution of *r* can affect the correlation time. If the relative change in variance was similar to the change in diffusion coefficient, the effects could approximately cancel, i.e. for compact proteins, moving in a narrower potential could compensate a slower diffusion coefficient. In fact, as might be anticipated from Fig. 5B, the variance does indeed decrease in the simulations for proteins with increasing FRET efficiency (*ρ*_MC_=-0.81 (0.05)), and similarly, the SAW − *ν* model predicts smaller variance as FRET efficiency increases (Fig. 5D). A direct experimental measure of the width of the distributions is provided by the variance of the FRET efficiency, Var[*E*], determined from a combination of ratiometric and lifetime-based FRET measurements.^56^ Indeed, the Var[*E*] from simulation and experiment are strongly correlated, with *ρ* = 0.88, confirming the observations from simulation. Although this is a simplified picture, a similar effect is expected in the more general case of diffusion within a potential well with position-dependent diffusion coefficients. An interplay between collapse and diffusion is also expected from models of polymer dynamics, such as the Rouse and Zimm models, where the chain relaxation time depends on the ratio of the chain dimensions and the diffusion coefficient determining the Brownian motion of the chain segments,^57,58^ and its role has been invoked to explain the invariance of the reconfiguration time of the charged IDP prothymosin *α* as a function of salt concentration. ^20^

While we have seen that the diffusion coefficients are not position-independent, they can be approximated as such, with the value of the effective constant diffusion coefficients being most sensitive to the diffusion coefficient near the peak in the distance distribution for each protein. If the diffusion model is refitted with the assumption of constant diffusion coefficients, we indeed find that the effective position-independent diffusion coefficient decreases with increasing FRET efficiency (*ρ*_MC_=-0.93 (0.02)), as expected (5E). The diffusion coefficient that would be inferred from the experimental *τ*_*r*_ and Var[*r*] from the SAW-*ν* model exhibits a very similar decrease. We can conduct a thought experiment to test the effect of these systematic changes in Var[*r*] and *D*_const_ by computing correlation times with Eq. 1 and assuming either (i) the same diffusion coefficient for all proteins (taken to be the average of those in Fig. 5E) or (ii) identical Var[*r*] for all proteins (taken to be the average of those in Fig. 5D). In each of these cases, there is a strong linear correlation of the calculated *τ*_*r*_ with ⟨*E*⟩, with correlation coefficients *ρ*_MC_ = −0.81 (0.05) and 0.88 (0.03) for cases (i) and (ii), respectively. Combining the effects of varying both Var[*r*] and *D*_const_ via Eq. 1 again yields almost no correlation with FRET efficiency (*ρ*_MC_ = 0.05 (0.12)), and in addition good agreement with *τ*_*r*_ computed from the full diffusion model (Fig. 5A). We can thus conclude that the small variation in *τ*_*r*_, and consequent insignificant correlation with FRET efficiency, is a consequence of almost perfectly compensating effects from the narrowing distribution of end-end distances for more compact IDRs and a local reduction in *D*(*r*) for shorter distances. We refer to these compensatory effects that help to maintain an almost invariant reconfiguration time as “dynamical buffering” in analogy to “conformational buffering”, compensatory changes in amino acid sequence composition and sequence length that have been suggested to lead to the conservation of optimal tethering in a large family of disordered adenovirus early gene 1A protein (E1A) linkers.^8^

Internal friction effects are also known to slow dynamics of unfolded and disordered proteins, particularly for more compact states^20,22^ – for example, the reconfiguration time of a small cold shock protein with approximately the same length as the proteins in the present work is over 100 ns in the unfolded state in the absence of denaturant, which can be attributed to its high internal friction. ^20^ Via analysis of the dependence of dynamics on solvent viscosity, we have determined that there is negligible internal friction in dCh−, one of the more expanded linkers (Fig. S7). The absence of internal friction in dCh− suggests low internal friction in the entire set of sequences. Based on the dynamic buffering effect we observe, the contribution of internal friction to the overall reconfiguration time cannot be greater than 10 to 15 ns, even for the most compact variants.

## Discussion

Although developments within the last decade have corrected the tendency for simulations of IDPs to give ensembles that were too collapsed, ^29,59^ and independent simulations by several groups have generally confirmed the accuracy of these new improvements. ^40,60–62^ However, it is also known that the properties of IDPs are strongly sequence-dependent, ^10,63–67^ a feature sometimes referred to as their “molecular grammar”. ^68,69^ Although it has been shown that force fields can reproduce the degree of compaction of selected proteins of different lengths,^60,61^ a more stringent test lies in a side-by-side comparison of proteins of the same length.

The functional versatility of IDPs comes from both the broad equilibrium ensemble of structures which they populate as well as how rapidly they explore it to find, form, and release interactions. These properties are exquisitely sensitive to the underlying sequence as well as the environment. The set of intrinsically disordered linker IDRs explored here showcases the diversity of ensembles that are populated, ranging from highly expanded sequences near the excluded volume limit to sequences that are slightly more compact than ideal chains, as characterized by single-molecule FRET spectroscopy. Since FRET measures properties averaged over a broad ensemble of structures, it is essential to complement such experiments with a theoretical or simulation model to aid in their interpretation. Analytical polymer models and coarse-grained molecular simulation models provide a starting point for analyzing structure and dynamics, ^10^ but all-atom simulations provide the highest spatial and temporal resolution and are able to predict both structural ensembles as well as absolute timescales of dynamics, provided the force fields used are of sufficient quality. However, it often is a challenge to sample long enough time scales for quantitative comparison to experiment.

In this work, we have successfully used long-timescale all-atom explicit-solvent simulations to characterize this set of 16 IDRs. Our simulations agree quantitatively with both single–molecule FRET efficiencies and, where available, SAXS, providing a consistent validation from complementary experiments. Further, by modeling the FRET dyes explicitly at atomistic detail, we find that the chromophores have a minimal influence on ensemble dimensions in these systems—consistent with the current view ^70,71^ and contrary to suggestions that common fluorophores collapse disordered chains.^49^ In fact, including dyes in the simulations yields closer correspondence to the measured FRET efficiencies without distorting the protein ensemble, in line with broader assessments of smFRET reliability.^31^ In contrast to the situation that would be expected if the dyes caused collapse, the FRET efficiencies including the explicit chromophores are systematically *lower* than without them. Taken together, this highlights that more rigorous methods of back-calculating experimental observables from simulations do result in better agreement with experiment.^31,72^

Nanosecond fluorescence correlation spectroscopy (nsFCS) combined with single–molecule FRET directly reports on end–to–end distance relaxation, or reconfiguration times, which are commonly on tens–of–nanosecond timescales for disordered proteins.^10^ By varying solution conditions and interpreting the results with polymer models, it has been possible to separate solvent friction effects from internal friction due to transient intrachain contacts. Extensive previous work has established that increased compaction tends to raise internal friction and slow intrachain motions. ^9,20–22^ Surprisingly, however, the reconfiguration times of the 16 linker IDRs reported here cluster around similar values and show no pronounced correlation with their large sequence–encoded differences in equilibrium compaction. All–atom explicit–solvent simulations reproduce this small variation in time scales together with the changes in compaction. The simulations can furthermore explain the apparent paradox with a simple mechanism: while more compact ensembles indeed exhibit reduced intrachain diffusivity, in accord with the established internal friction picture, the narrower end-to-end distribution for those proteins has the opposite effect. Because reconfiguration times can be approximated as 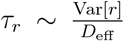 the decrease in variance offsets the decrease in effective diffusivity, preserving a nearly constant *τ*_*r*_ across sequences.

Note that this observed compensation may not extend to scenarios of more extreme compaction approaching the globule limit, where internal friction effects are felt most strongly, and a sharper reduction in diffusion coefficient may be expected. Indeed, simulations of a coarse-grained model with explicit solvent also showed a dynamical buffering effect, with almost constant reconfiguration times over a range of solvent quality and degree of chain collapse, and a sharp slowdown only being observed once the chain approaches the most compact extreme as *ν* → 1*/*3.^27^

The quantitative description of both equilibrium structure and dynamics in disordered protein ensembles which we present here has been enabled by progressive refinement of all–atom MD force fields over the years so as to correct systematic biases that hampered earlier simulations.^18^ Most notably, secondary structure biases that favored either *α*-helical or *β*-sheet structure^73^ and the tendency for unfolded proteins to be too collapsed^74^ were obviously detrimental for simulations of disordered proteins. Guided by empirical data, however, it was possible to improve simulations so as to match experiment.^29,59,75^ The present set of 16 disordered linker IDRs presents a particularly stringent test of these improvements, as all are of the same length, and thus only sequence effects, ^10,63–67^ sometimes referred to as “molecular grammar”^68,69^ are responsible for the differences between them.

The good agreement of the simulations with experiment, both in terms of overall dimensions and reconfiguration times strongly validates the force field improvements. Nonetheless, analysis of residual deviations from experiment suggests that there are some discrepancies outside of the statistical noise, and these appear to be most highly correlated with the presence of oppositely charged residues. Simple reweighting procedures adding a correction term for salt bridge formation suggest that salt bridges, particularly those involving arginine, are slightly too strong in the force field, providing an avenue for future force field improvement.

The insights obtained here into the dynamics of disordered linkers may also have implications for their biological function. In signaling and regulation, preserving fast chain reconfiguration regardless of average compaction ought to facilitate conformational search, encounter–complex formation, and rebinding.^1^ For example, in high–affinity disordered or “fuzzy” complexes, polyelectrolyte interactions can enable strong binding with rapid on/off kinetics, precisely the context where maintaining rapid reconfiguration is advantageous.^51,52^ The dynamical buffering which conserves these rapid timescales even as configurational properties are modulated by sequence, also allows for robustness during evolution, reminiscent of the “conformational buffering” role proposed for the sequences of disordered linker regions.^8^

## Methods

### All-atom simulations

Extended all-atom configurations of all 16 IDR variants without dyes were generated using *tleap* from the Amber package.^76^ The resulting structures were then placed into 14-nm rhombic dodecahedral boxes. To prevent self-interaction across the periodic boundaries, short vacuum simulations were carried out for each IDR, and conformations with a maximum end-to-end distance below 12.5 nm were retained. Finally, for each of the 16 IDRs, three of these conformations were randomly chosen to initiate all-atom explicit-solvent simulations. The selected structures were energy-minimized using the steepest-descent algorithm. Each simulation box was then solvated with the TIP4P2005s water, ^29^ and the systems were again energy-minimized using the steepest-descent algorithm. Subsequently, the desired number of potassium and chloride ions were inserted to obtain a 172 mM KCl salt concentrations, matching the total ionic strength of the KCl salt and buffer used in the experiment. Subsequently, the system was again energy-minimized. The total number of atoms per simulation was approximately 250,000, varying slightly among different proteins. All simulations were performed using GROMACS^77^ version 2021.5. Protein interactions were modeled using Amber99SBws, ^29^ with protein-dye and dye-dye interaction parameters described previously^30^ and using optimized dye-water interaction parameters in which dye-water interactions were scaled by a factor 1.15 – this was shown to result in improved dynamical properties for AlexaFluor chromophores^31^ and the same factor was used here for Cy3B and CF660R. The integration time step was 2 fs. The temperature was kept constant at 295.15 K using velocity rescaling^78^ (*τ* = 1 ps), and the pressure was kept at 1 bar using the Parrinello-Rahman barostat^79^ (*τ* = 5 ps). Long-range electrostatic interactions were modeled using the particle-mesh Ewald method.^80^ Dispersion interactions and short-range repulsion were described by a Lennard-Jones potential with a cutoff at 1 nm. H-bond lengths were constrained using the LINCS algorithm.^81^ Each independent run was at least 2.05 *µ*s, and the first 50 ns were treated as equilibration and were omitted from the analysis.

For all 16 IDRs, the protein coordinates from each of the three independent runs without dyes were extracted after 100 ns of simulation time and used as starting structures for simulations with dyes. The explicit Cy3B and CF660R dyes were attached using custom Python scripts. All subsequent steps—including energy minimization, solvation with TIP4P2005s water, and addition of potassium and chloride ions to reach 172 mM KCl—were identical to those used for the simulations without dyes. Each independent run was at least 2.02 *µ*s long, with the first 20 ns treated as equilibration and omitted from the analysis.

The mean transfer efficiency, ⟨*E*⟩, was obtained from simulations with dyes as described previously.^30,82^ In short, we computed the survival probability of the donor excited state with a time-dependent transfer rate and averaged it over all possible excitation times *t*_0_ along each trajectory:

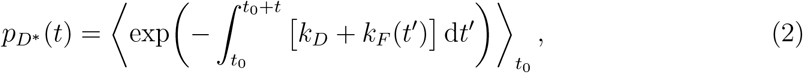

where

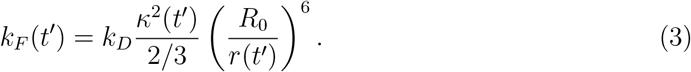

Here, *κ*^2^ is the orientational factor for the donor–acceptor dye pair, *r* is the instantaneous inter-dye distance, and *R*_0_ is the Förster radius for *κ*^2^ = 2*/*3. The dyes were treated as non-emissive whenever any donor–acceptor atom pair was closer than 0.4 nm (van-der-Waals contact). The average FRET efficiency was then computed by integrating the survival probability:

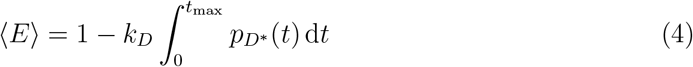

with *t*_max_ = 20 ns, chosen in order that that 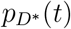 had effectively decayed to zero. From the simulations without dyes, the inter-dye distance *r* was estimated as

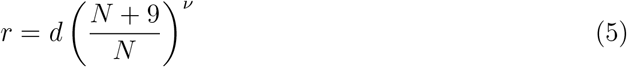

where *d* is the C*α*–C*α* distance between the labeled residues. Here, *N* is the sequence separation between the labeling sites, and we used a scaling exponent *ν* = 0.6. This approach accounts for the dye and linker lengths by adding nine “effective” residues to *N*.^48^ The estimate is only weakly sensitive to *ν*: varying *ν* by ±0.1 changes the inferred transfer efficiency by roughly ±0.01. For transfer efficiency calculations from simulations without explicit FRET dyes, we assumed *κ*^2^ = 2*/*3.

The number of possible salt bridges (Fig. 3) was computed as the number of oppositely charged residue pairs within each IDR sequence, considering only pairs separated by at least three residues along the sequence. Residue–residue contact fractions (Fig. S5) were evaluated using a transition/core-state approach.^83^ Instead of a single cutoff separating bound and unbound states, distinct thresholds were applied for contact formation and breaking. For each residue pair, the distance metric was the minimum heavy-atom distance between the two residues. Starting from an unformed state, a contact was considered formed when this distance dropped below 0.38 nm; once formed, it remained in the contact state until the distance exceeded 0.8 nm.^83^

### Force field parameters for Cy3B and CF660R

Parameters for the chromophores Cy3B and CF660R, and their cysteine-conjugated linker, were derived using the Antechamber program from the AmberTools package.^84^ The labelled residue was split into a part corresponding to the linker-conjugated cysteine, and a second part corresponding to either the Cy3B or CF660R chromophore. Following the nomenclature of the AMBER-DYES force field, ^85^ the internal Cysteine and linker were denoted “C2R”, while Cy3B and CF660R were respectively named “C3B” and “CF6”. Because the C2R residue in the AMBER-DYES force field is much more polarized relative to the standard AMBER94 cysteine, ^86^ and this is known to be correlated with aberrant secondary structure formation, ^87^ we derived new charges for the C2R residue by constraining the backbone charges to be the same as the others in AMBER94-derived force fields, basing the calculations on surface electrostatic potentials determined with Gaussian 16 using Hartree-Fock with the 6-31G* basis set. The chromophore parameters were obtained using a similar process to those for C2R, constraining the charges of the amide conjugation to the linker to be the same as for the protein backbone, but using a 6-31+G* basis set (this having previously given better results for AlexaFluor dyes ^30^). Scripts used to generate the new parameters and the parameters themselves are available on the accompanying Zenodo repository (https://doi.org/10.5281/zenodo.13381447), as is the script for generating initial structures of the dye-conjugated proteins.

### Reweighting to estimate the effect of salt bridge strength

We assume that the the true energy of each salt bridge, considered for simplicity as a contact energy, is *E*_sb_ = *E*_sb,ff_ + *ϵ*_sb_, where *E*_sb,ff_ is the energy of the salt bridge resulting from the force field used. Assuming all other terms in the energy function are unchanged, the total difference in energy between the correct salt bridge energy and the force field salt bridge energy would be Δ*E* = *N*_sb_*ϵ*_sb_ (in units of *k*_B_*T*). Each frame *i* of the simulation, having *N*_sb_ salt bridges, can then be reweighted using weight *w*_*i*_ ∝ exp[−*N*_sb_(*i*)*ϵ*_sb_], allowing observables such as FRET efficiency to be computed with the correction. A simple scan over values of *ϵ*_sb_ identifies the optimal value as that which best matches the reweighted simulation efficiencies to experiment. We also considered the case where specific corrections are made for each of the four types of salt bridge (Glu-Lys, Glu-Arg, Asp-Lys, Asp-Arg), i.e. Δ*E* = *N*_EK_*ϵ*_EK_ + *N*_ER_*ϵ*_ER_ + *N*_DK_*ϵ*_DK_ + *N*_DR_*ϵ*_DR_. In this case the optimal parameters were determined by gradient descent using an adaptive step size.^88^ The analytical gradients of *χ*^2^ for each protein with respect to *ϵ*_AB_ are given by

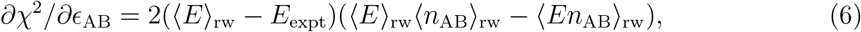

where the average ⟨·⟩_rw_ indicates the reweighted average using the current contact energy parameters, and *E*_expt_ is the experimentally observed value. The total gradient is the sum of Eq. 6 over all the proteins. In practice, we minimized the target function *L* = *χ*^2^ + ^1^ *γ*(*ϵ*_EK_ + *ϵ*_ER_ + *ϵ*_DK_ + *ϵ*_DR_)^2^, where the constant *γ*, chosen to be 1.0, penalizes large values of the contact energies.

### Bayesian reweighting

Simulations with dyes were reweighted using Bayesian inference of ensembles (BioEn)^43^ to reach agreement with both the means and the variances of the transfer efficiency distributions observed experimentally. It is worth emphasizing, as already discussed in Ref. 46, that we use for this approach not the variance of the transfer efficiency histogram, but the variance of the transfer efficiency distribution that corresponds to the underlying distance distribution, which can be obtained from the deviations of the mean fluorescence lifetimes from the static FRET line.^56,89,90^ In this way, the reweighting takes into account not only experimental information on the average intramolecular distance but also on the variance of the distance distribution. The instantaneous transfer efficiency was determined from simulations using the dye coordinates every 20 ps. The uncertainty of the experimental transfer efficiency was set to 0.03 and the uncertainty of the variance of the transfer efficiency distribution was set to 0.02.^46^ Reweighting parameters and results are shown in Table S3. Histograms of the initial and reweighted distance distributions are shown in Fig. S3.

### SAXS calculations

To calculate SAXS curves (Fig. 2B), we saved protein structures every 5 ns from all runs of all simulations (1200 PDB files per IDR). Subsequently, we used CRYSOL^91^ with default parameters to compute SAXS curves for each structure. We note that the experimental curve was not provided to CRYSOL, to avoid any overfitting that could arise from adjusting the hydration-layer density to better match the experimental SAXS profile.^92^ For each IDR, the 1200 calculated SAXS curves were then averaged, and the error at every value of momentum transfer, *q*, was determined as the standard deviation across the set of curves. To overlay experimental and computed SAXS curves, we scaled only the absolute intensity of the experimental curve, without applying additional corrections for possible buffer mismatch.^93^ We note that explicit consideration of the solvent environment typically yields protein scattering curves very similar to those from CRYSOL, likely because the solvation shell is highly correlated with the path of the protein chain. ^94^ Experimental SAXS curves have been taken from the previous study. ^19^

### Computing reconfiguration times

Dynamics of the interchromophore distance was analyzed with a one-dimensional diffusion model.^41,54^ Briefly, the distance coordinate was discretized into 30 equal-size, non-overlapping bins between the minimum and maximum sampled value for each trajectory. Histograms of the number of observed transitions between bin *i* at time *t* and bin *j* at time *t* +Δ*t, N*_*ji*_(Δ*t*), were determined for different lag times Δ*t*. Discretized free energies and position-dependent diffusion coefficients were optimized via Monte Carlo simulations using the log-likelihood function

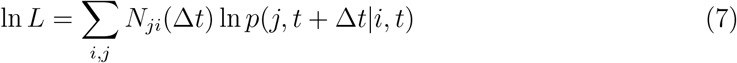

where the propagators *p*(*j, t* + Δ*t*|*i, t*) describing the conditional probability of being in bin *j* a time Δ*t* after having been in bin *i* are obtained from the discretized diffusion model as previously described:^41,54^ in short, the discretized dynamics is mapped to a chemical kinetics scheme describing the time evolution of populations in the bins, 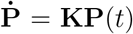 where **P**(*t*) is the vector of the bin populations at time *t*, and **K** is a rate matrix derived from the diffusion coefficient(s) *D*_*i*_ and free energies *F*_*i*_ associated with each bin according to the scheme of Bicout and Szabo.^54,95^ The propagators are then given by *p*(*j, t* + Δ*t*|*i, t*) = (exp[Δ*t***K**])_*ji*_. In estimating the most probable parameters from the data, a uniform prior is used for the diffusivities and free energies. We can compute normalized correlation functions

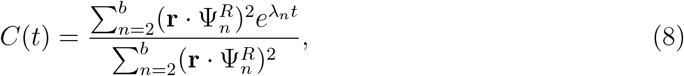

where the elements of **r** are the centers of each bin on the distance coordinate, 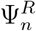 is the *n*th right eigenvector of **K** (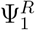 is the stationary eigenvector), and *λ*_*n*_ is the *n*th eigenvalue. From this, the correlation times follow:

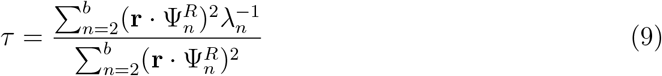

In determining the optimal parameters, we fitted simultaneously to lag times of 5, 10 and 20 ns, so as to include both short and long time correlations; the convergence of the relaxation times with respect to the lag time for fits to single lag times is shown in Fig. S8.

Correlation coefficients *ρ*_MC_ and associated errors were computed by generating 100 synthetic, random data sets using the mean and standard error for each quantity. For each data set, the linear correlation coefficient *ρ* was determined, and *ρ*_MC_ and its associated error were determined from the mean and standard deviation of the correlation coefficients of individual synthetic data sets.

### Single-molecule fluorescence spectroscopy

The set of 16 naturally occurring IDRs, each comprising 57 amino acid residues, was selected from the linker regions connecting folded domains in RNA-binding proteins, as described previously.^19^ For fluorescence labeling, cysteine residues were introduced at the the N- and C-termini of each IDR sequence. The constructs were recombinantly expressed in *Escherichia coli*, purified, and subsequently labeled with Cy3B and CF660R fluorophores (Förster radius 6.0 nm) via maleimide chemistry as described previously.^19^ For single-molecule experiments and nanosecond fluorescence correlation spectroscopy (nsFCS),^10,13^ donor–acceptor–labeled peptides were diluted in 20 mM KH_2_PO_4_/K_2_HPO_4_, 125 mM KCl, pH 7.3, supplemented with 0.001% Tween-20 and 10 mM DTT. Measurements were performed at 22 ^*°*^C either in chambered cover slides (*µ*-Slide, ibidi) at roughly 0.1 nM protein concentrations or, if possible, in zero-mode waveguides (ZMW) at high nanomolar concentrations to reduce acquisition times.^46,72^

Measurements were carried out either on a MicroTime 200 (PicoQuant) or on a custom-built confocal single-molecule instrument. Excitation lasers were operated in continuous-wave mode. The MicroTime 200 was equipped with an Olympus UplanApo 60×/1.20 W objective. Donor excitation was achieved using an Oxxius laser at 532 nm attenuated to 50 *µ*W at the back aperture of the objective. Fluorescence was collected through the same objective and separated from backscattered light by a triple-band mirror (ZT405/530/630RPC, Chroma Technology). Residual excitation light was further suppressed with a long-pass filter (LP532, Chroma Technology) before passing through a 100 *µ*m pinhole. Emission was split into four detection channels using a 50/50 or polarizing beam splitter and two dichroic mirrors (635DCXR, Chroma Technology). Donor and acceptor signals were subsequently filtered with an ET585/65M band-pass (Chroma Technology) and LP647RU long-pass filter (Chroma Technology) / 750 nm blocking edge BrightLine multiphoton short-pass emission filter (Semrock), respectively, and detected by four single-photon avalanche diodes (SPCM-AQR-15, PerkinElmer Optoelectronics). Photon arrival times were recorded with 16 ps resolution using a HydraHarp 400 TCSPC module (PicoQuant). In the custom-built instrument, the excitation source and optical filters were identical to those in the commercial instrument.^96^

For data analysis, we used Fretica, a Wolfram Mathematica package with a backend written in C++ available from https://github.com/SchulerLab/Fretica. Only photons from bursts of molecules double-labeled with active donor and acceptor fluorophores were used for nsFCS analysis, which reduces the contribution of donor-only and acceptor-only signal to the correlation. The selection was based on the transfer efficiency. All experimental transfer efficiencies reported here are from Holla et al. ^19^ The correlation between two time-dependent intensity signals, *I*_*a*_(*t*) and *I*_*b*_(*t*), recorded on two detectors, *a* and *b*, is defined as

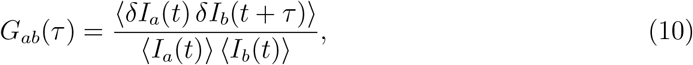

where the angle brackets denote time averaging. In our experiments, two donor detection channels and two acceptor detection channels were used, yielding the autocorrelation functions *G*_*dd*_(*τ*) and *G*_*aa*_(*τ*). In addition, photons detected in both donor and acceptor channels were used to calculate the cross-correlation function *G*_*da*_(*τ*). The resulting nsFCS curves were computed and fitted with *τ*_*cd*_ as a global parameter over a linear range of lag times from 0 ns to 100 ns using the fit equation

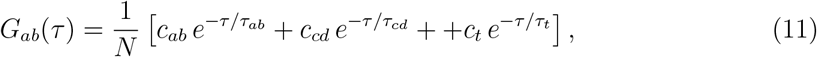

where *N* is a normalization factor. The first term, with amplitude *c*_*ab*_ and characteristic timescale *τ*_*ab*_, describes photon antibunching, the second term, with amplitude *c*_*cd*_ and timescale *τ*_*cd*_, describes chain dynamics, and the third term, with amplitude *c*_*t*_ and timescale *τ*_*t*_, describes triplet blinking.

We analyzed the correlation times corresponding to chain dynamics as described previously,^46,58^ where the distance dynamics are treated as diffusive motion in the potential of mean force associated with the distance distribution. In this framework, for any distance-dependent observable, *X*, the correlation time, *τ*_*X*_, is defined as

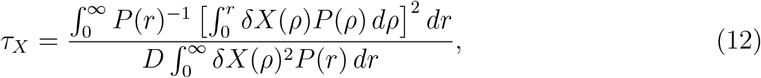

where *D* is the effective dye-to-dye diffusion coefficient, and *δX*(*r*) = *X*(*r*) − ⟨*X*(*r*)⟩_*r*_. The experimentally measured intensity correlation time from nsFCS, *τ*_*cd*_ = *τ*_*E*_, corresponds to the observable *X* = *E*, the transfer efficiency. To obtain the chain reconfiguration time, *τ*_*r*_, which corresponds to correlation time of the dye-to-dye distance, *X* = *r*, we computed the conversion factor *ϑ* = *τ*_*r*_*/τ*_*E*_ by evaluating the general expression for the correlation time *τ*_*X*_ with the distance distribution approximating a self-avoiding walk model of a polymer chain, characterized by the length scaling exponent *ν*.^23^ *ϑ*, which depends on the chosen distance distribution, the measured transfer efficiency, *E*, and the Förster Radius, *R*_0_ (here 6.0 nm), was then applied to convert the measured *τ*_*E*_ into *τ*_*r*_. The total uncertainty of the measured fluorescence decay times was obtained by combining statistical and systematic contributions in quadrature. The systematic uncertainty was estimated from the fitting procedure of the nsFCS data by varying the time window used for the fit from 100 to 1000 ns and using the standard deviation of the results. The statistical uncertainty was estimated from replicate measurements available for three of the samples, yielding a value of 10%.

In addition to the nsFCS analysis, FRET efficiency correlation analysis was performed, as described by Terterov et al.^97^ The correlation curves were fitted with

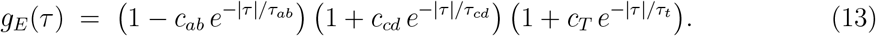

The three terms with amplitudes *c*_*ab*_, *c*_*cd*_, *c*_*t*_, and timescales *τ*_*ab*_, *τ*_*cd*_, *τ*_*t*_ describe photon antibunching, chain dynamics, and triplet blinking, respectively. Estimates of systematic uncertainty of the FRET efficiency correlation fits were obtained using a two-step fitting approach based on Eq. (13). First, each protein variant was fitted individually. In a second step, all data sets were fitted globally using Eq. (13) with *τ*_*t*_ as a shared parameter. The standard deviation of the resulting values of *τ*_*cd*_ from the individual and global fits is reported as the systematic uncertainty. The statistical uncertainty was again estimated from the analysis of replicate measurements available for three of the samples. The reconfiguration times reported here (Figures 2 and 5) were calculated as the arithmetic averages of the values from the nsFCS and the FRET efficiency correlation analysis. To be able to report a single value of the uncertainty for each protein variant in the figures, the total uncertainty for the reconfiguration time is shown by adding the statistical and systematic components from both methods in quadrature as:

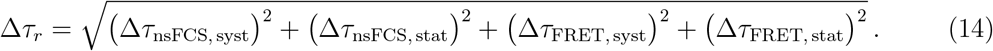

All reconfiguration times and the individual components of the associated uncertainties are reported in Table S2.

Viscosity-dependent nsFCS measurements of dCh− were performed using ZMWs as described above. Measurements were carried out in 20 mM potassium phosphate buffer, 125 mM KCl, pH 7.3, 0.001% Tween-20, and 10 mM DTT, with 0, 2, 5, 10, and 20% (v/v) glycerol. Solution viscosities were calculated from measured refractive indices and based on reference values from the CRC Handbook of Chemistry and Physics (63rd edition). Correlation curves were fitted over a 0.1 *µ*s window without a triplet term and over a 0.5 *µ*s window including a triplet term. The mean of the resulting decorrelation times was used to calculate the reconfiguration times as described above. An uncertainty in the reconfiguration times of 10%, as estimated for the other nsFCS measurements, was assumed.

## Acknowledgement

We thank Daniel Nettels for discussion and help with optimizing data analysis tools, Steffen Winkler and Vincent Maximilian Münch for help with sample preparation and measurements, and Kresten Lindorff-Larsen for discussions based on Martini simulations. RB was supported by the Intramural Research Program of the National Institute of Diabetes and Digestive and Kidney Diseases (NIDDK) within the National Institutes of Health (NIH). The contributions of the NIH author are considered Works of the United States Government. The findings and conclusions presented in this paper are those of the author(s) and do not necessarily reflect the views of the NIH or the U.S. Department of Health and Human Services. BS was supported by the Swiss National Science Foundation (Grant numbers 310030 197776, CRSII5 205922). We used the computational resources of Piz Daint, Alps, and Eiger at the CSCS Swiss National Supercomputing Center, and of the National Institutes of Health HPC Biowulf cluster (http://hpc.nih.gov).

## Supporting Information Available

A supplementary file containing seven figures and three tables is provided with the manuscript.

## TOC Graphic

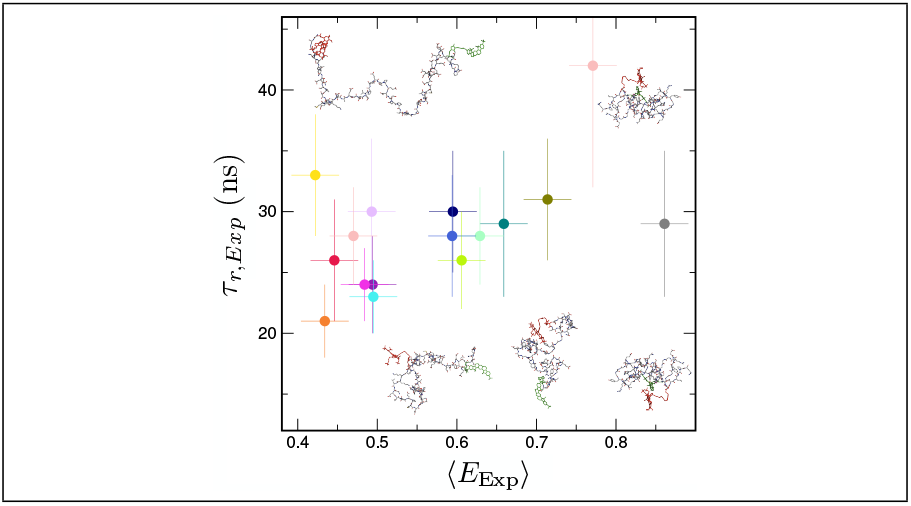

