## Supplementary Information for "Dynamical buffering of reconfiguration dynamics in intrinsically disordered proteins"

### Supporting Figures

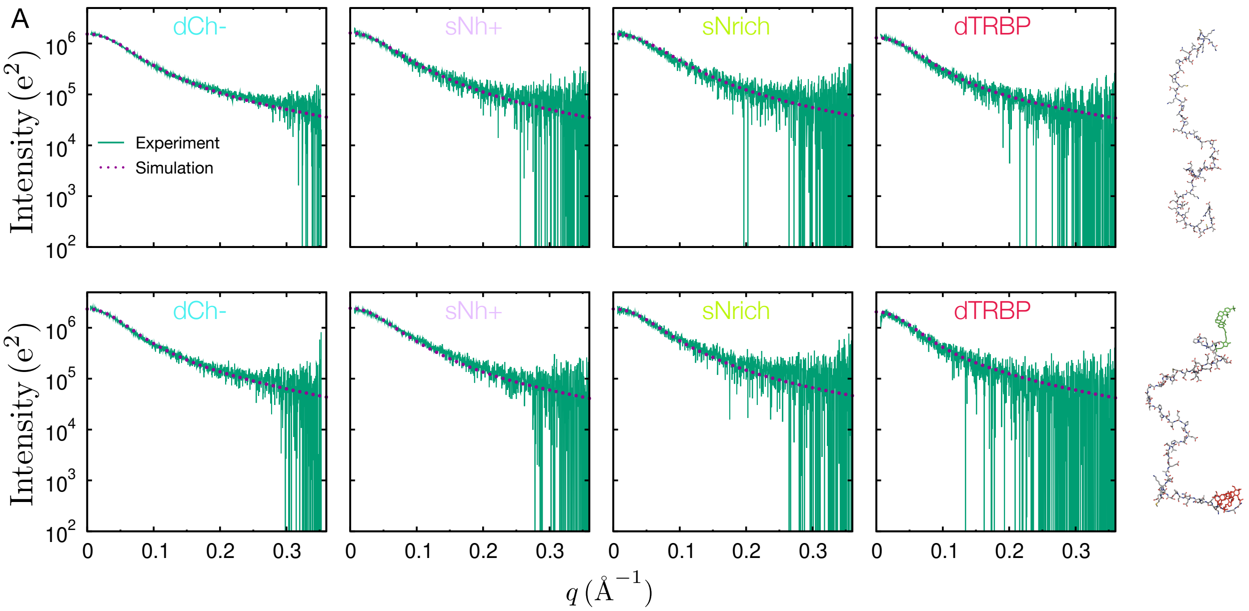


**Figure S1:** Comparison of the experimental SAXS curves with those calculated from the simulations. (A) without dyes; (B) with dyes (note: in the experiment, IDRs were double-labeled with Alexa 488,[1] while in the simulation, Cy3B + CFR660R were used).

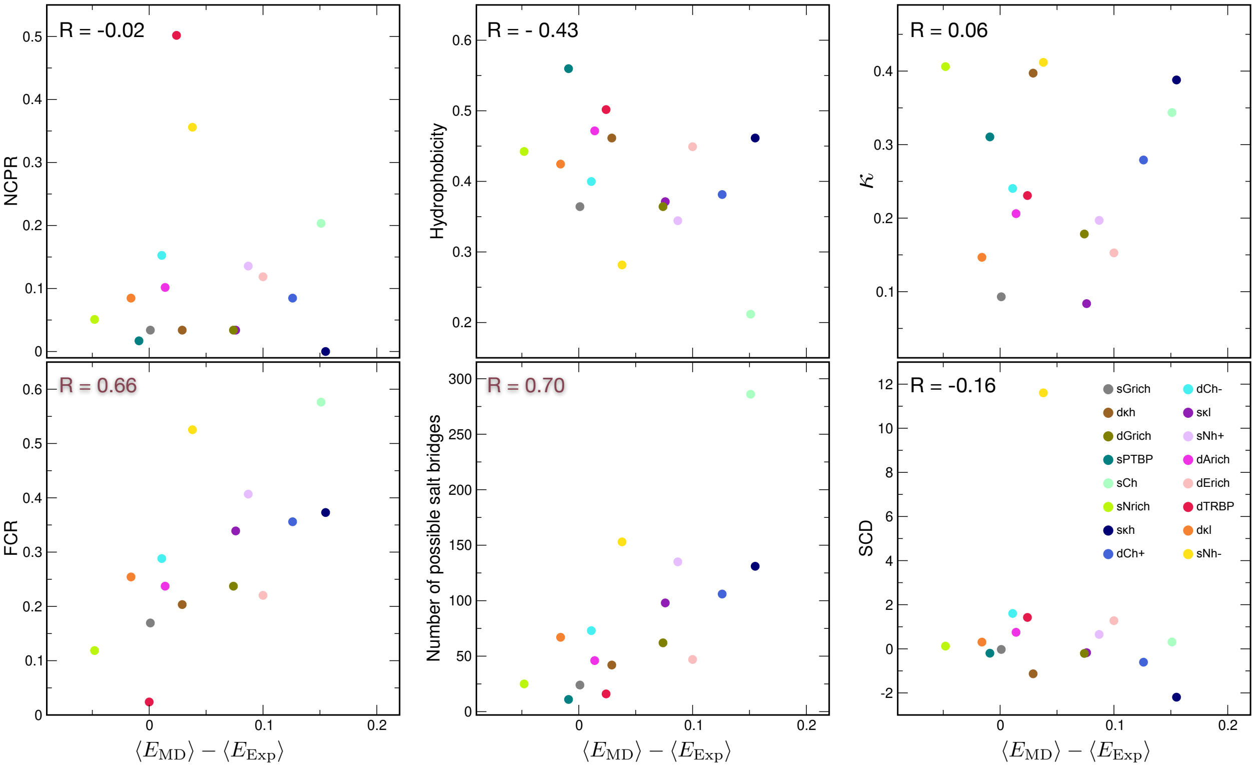


**Figure S2:** Correlations between the deviation of simulated and experimental FRET efficiencies and net charge per residue (NCPR), fraction of charged residues (FCR), hydrophobicity, number of possible salt bridges, $\kappa$ parameter[2] and sequence charge decoration (SCD).[3]

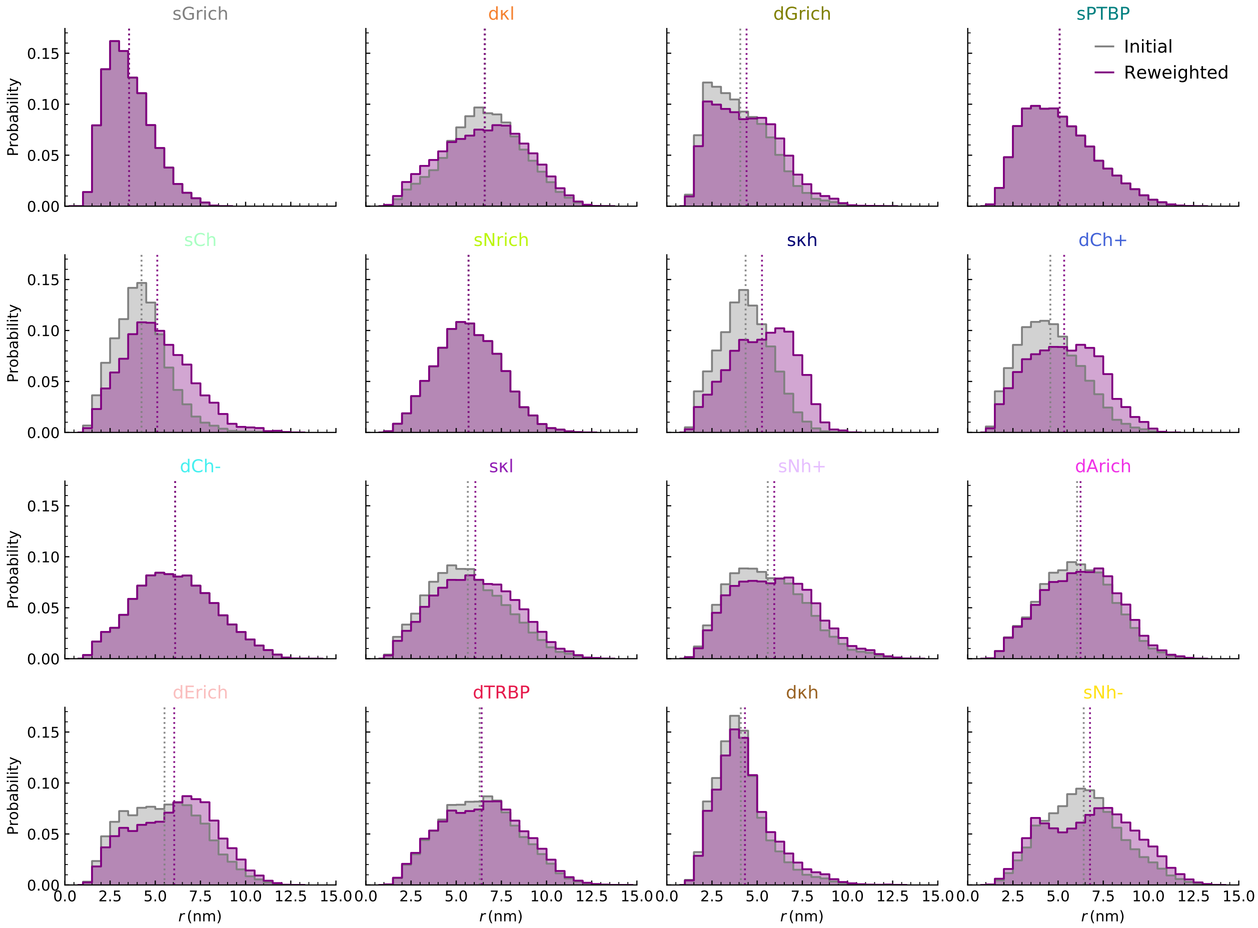


**Figure S3:** Dye-dye distance distributions before and after reweighting. The vertical lines represent the average values.

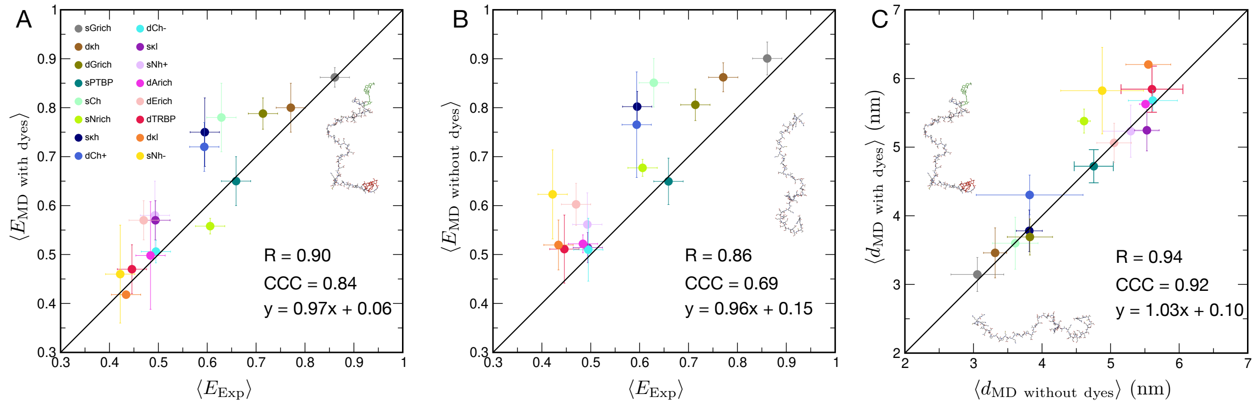


**Figure S4:** Comparison of experimental transfer efficiencies and transfer efficiencies calculated from simulations with (A) and without (B) dyes. Transfer efficiencies from simulations with dyes are slightly lower than those from simulations without dyes (see also Fig. 4A), suggesting slightly more extended IDRs when dyes are included. However, Cys–Cys distances from the two simulation sets (C) indicate that average IDR dimensions are essentially identical with and without dyes (see also Fig. 4B). Taken together, these results indicate that the dyes do not appreciably perturb the conformational ensemble of the IDRs; the small differences in transfer efficiencies likely arise from the dyes' excluded volume and slightly anisotropic positioning in the local protein environment (see Fig. S5). R denotes the Pearson correlation coefficient and CCC the concordance correlation coefficient.

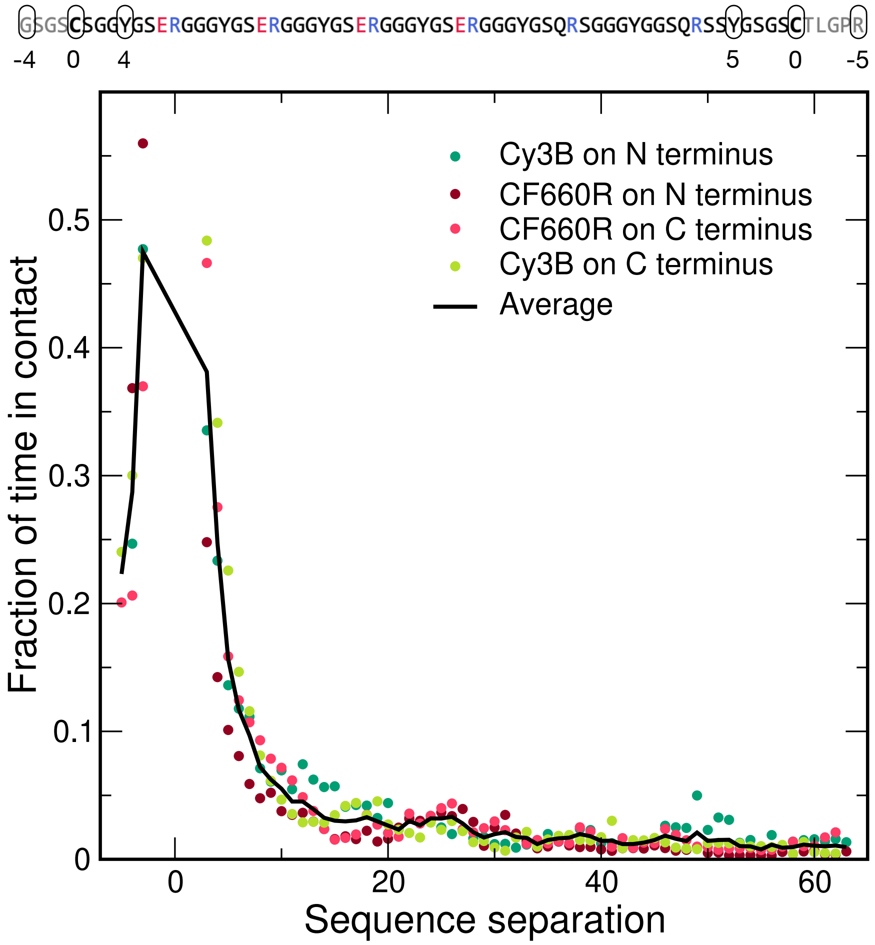


**Figure S5:** Fraction of time (normalized per simulation length) that the dyes are in contact with protein residues as a function of sequence separation. Negative sequence separation values indicate residues located closer to the termini than the Cys positions used for labeling (see schematic above the plot). The dyes predominantly contact neighboring residues, with slightly longer interaction times observed near the termini. This trend is consistent with the slightly reduced transfer efficiencies in simulations with dyes compared to those without dyes (Fig. 4 and Fig. S4). For details on the calculation of the fraction of time in contact, see Methods.

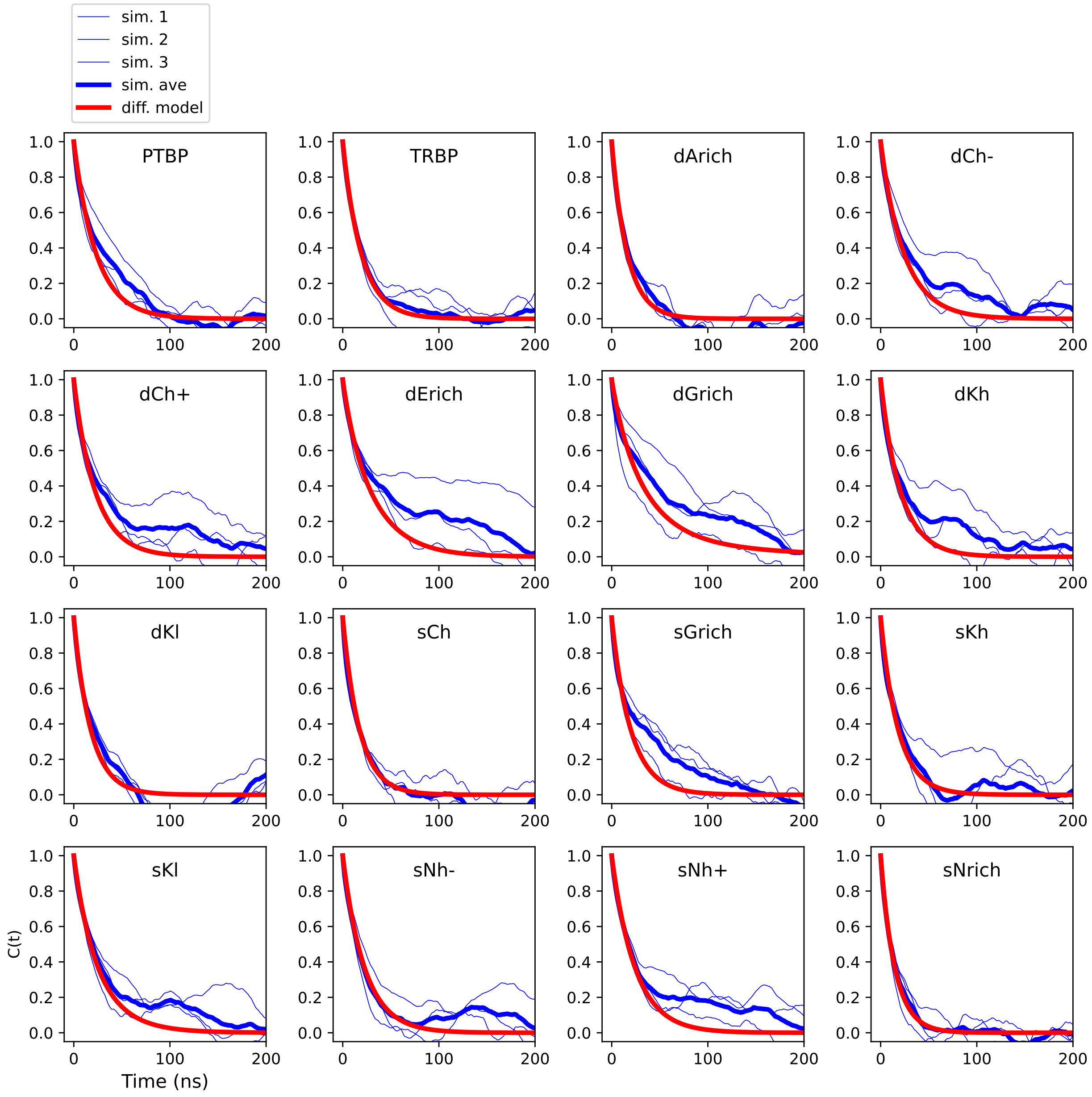


**Figure S6:** End-to-end distance correlation functions from the simulations. For each linker IDR, the correlation functions are plotted for each simulation replicate individually (thin blue lines), for the average over the three replicates (thick blue lines), and for the prediction using the one-dimensional diffusion model (thick red lines).

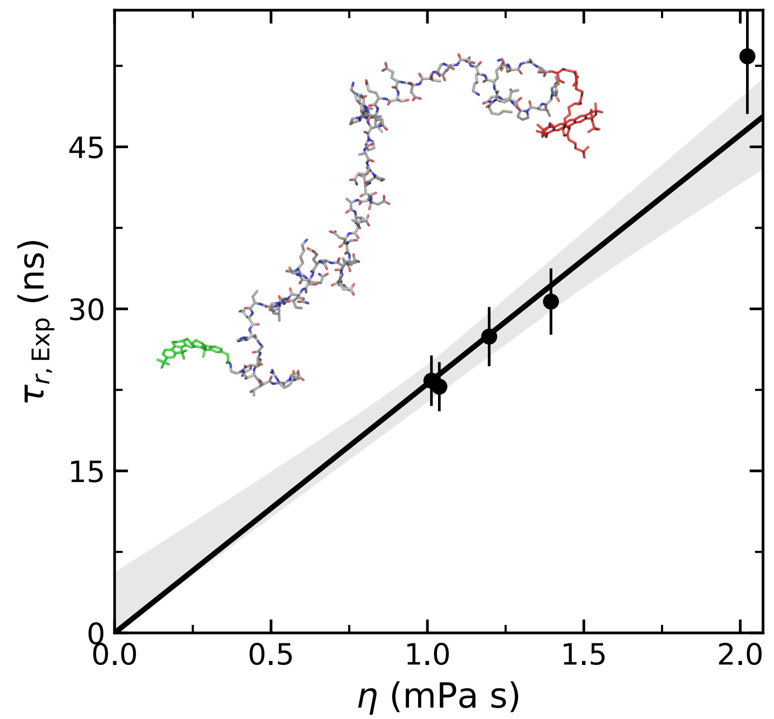


**Figure S7:** Quantifying internal friction in dCh− from solvent viscosity-dependent chain reconfiguration times. The internal friction time,[4] $\tau_i$, results from extrapolating the measured reconfiguration times, $\tau_r$, to zero solvent viscosity. Here the fit was constrained to yield $\tau_i \geq 0$, since internal friction times cannot take negative values. Inset: representative snapshot of dCh−.

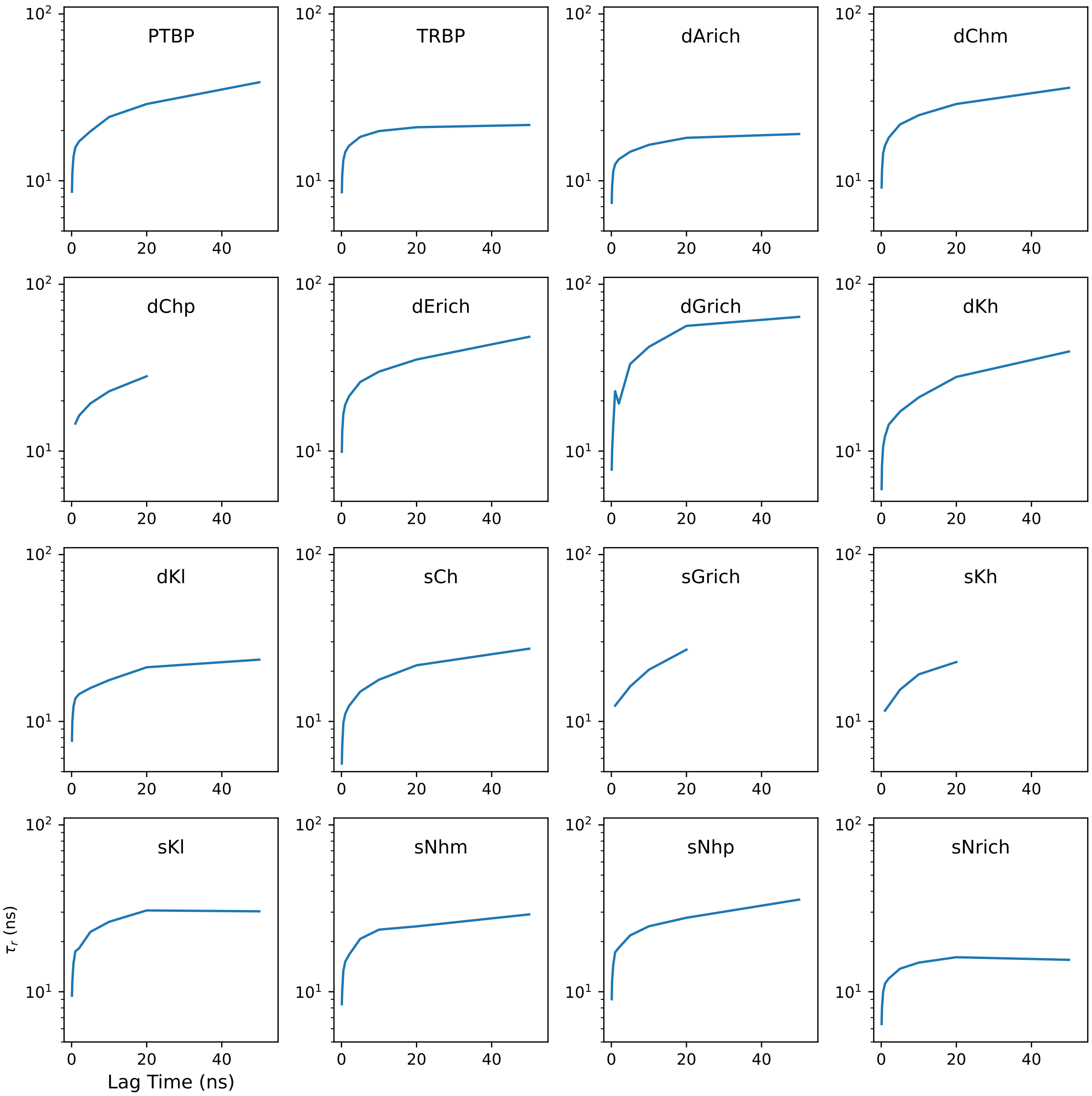


**Figure S8:** Dependence of reconfiguration time inferred from diffusion model on lag time (using data from single lag to fit model).

### Supporting Tables

Table S1: Sequences of all linker IDRs used in simulations and experiments. Donor-/acceptor-labeling at the termini was site-selective.[1] R denotes the red dye (CFR660R, net charge $Q = -1$); G denotes the green dye (Cy3B, net charge $Q = 0$). Cys residues (positions 5 and 63) were used for labelling. 'GR' labelling refers to Cy3B at position 5 and CF660R at position 63, while 'RG' indicates the opposite.

| Linker | Sequence | Site-specific labelling |
|---|---|---|
| sGrich | GSGSCSGGYGSERGGGYGSERGGGYGSERGGGYGSERGGGYGSQRSGGGYGGSQRSSYGSGSCTLGPR | GR |
| d$\kappa$h | GSGSCAMGGGPGPGTDFTSDQADFPDTLFQEFEPPAPRPGLAGGRPGDAALLSAAYGRRRLLCTLGPR | GR |
| dGrich | GSGSCGPRTGLEGAGMAGGSGQQKRVFDGQSGPQDLGEAYRPLNHDGGDGGNRYSVIDRIQECTLGPR | GR |
| sPTBP | GSGSCRPDLPSGDSQPSLDQTMAAAFGLSVPNVHGALAPLAIPSAAAAAAAAGRIAIPGLAGCTLGPR | RG |
| sCh | GSGSCKKAQPEMNDKDDNESGNEDAEENHDDEEDENEEEDRQVDQASKNKESKRKAQNKREDCTLGPR | GR |
| sNrich | GSGSCLDQEDNNGPLLIKTANNLIQNNSNMLPLNALHNAPPMHLNEGGISNMRVNDSLPSNTCTLGPR | GR |
| s$\kappa$h | GSGSCKPRRLSKLRRSKKPADEENNAASQDPTVEATQERGQASEDPENAANNAKQAKPTSDDCTLGPR | RG |
| dCh+ | GSGSCQTPLKRIKVKTPGKSGAAAREGSVVSGTDGPTQTGKPERRKRLNPPKDKLIDMDDADCTLGPR | RG |
| dCh− | GSGSCMGLPTGMEEKEEGTDESEQKPVVQTPAQPDDSAEVDSAALDQAESAKQQGPILTKHGCTLGPR | RG |
| s$\kappa$l | GSGSCAPEGVFKLPAPPKEIKKSEKSEKSSDSINDKKEVESTTKTAATTTTTKKGTDNNTQFCTLGPR | RG |
| sNh+ | GSGSCPEEIETRKKDRKNRREMLKRTSALQPAAPKPTHKKPVPKRNVGAERKSTINEDLLPPCTLGPR | GR |
| dArich | GSGSCLPTGAEGRDSSKGEDSAEETEAKPAVVAPAPVVEAVSTPSAAFPSDATAEQGPILTKCTLGPR | RG |
| dErich | GSGSCEGEMFGVMSEENSVVLSVEQPAELKEVADVSPPPTTRNHTIEMKPPLSAQQSESNPVCTLGPR | RG |
| dTRBP | GSGSCLEPALEDSSSFSPLDSSLPEDIPVFTAAAAATPVPSVVLTRSPPMELQPPVSPQQSECTLGPR | RG |
| d$\kappa$l | GSGSCLPTGMDQKTTELMKVDEPVSTQETLPPVIKMEPEPVPITETPTDENARQQGPILTKHCTLGPR | GR |
| sNh− | GSGSCKKKLQDTPSEPMEKDPAEPETVPDGETPEDENPTEEGADNSSAKMEEEEEEEEEEEEECTLGPR | RG |

Table S2: Reconfiguration times from nsFCS and FRET-corrected analyses. The reconfiguration times for each method are denoted $\tau_{\mathrm{nsFCS}}$ and $\tau_{\mathrm{FRET}}$; their statistical uncertainties are fixed at 10% of the corresponding value (determined from repeat measurements; see Methods). The combined value $\tau_r$ is the arithmetic mean of $\tau_{\mathrm{nsFCS}}$ and $\tau_{\mathrm{FRET}}$. The uncertainty $\Delta\tau_r$ is the quadratic combination of the statistical and systematic uncertainties from both methods.

| Linker | $\tau_{\mathrm{nsFCS}}$ (ns) | $\Delta\tau_{\mathrm{nsFCS,\,syst}}$ (ns) | $\tau_{\mathrm{FRET}}$ (ns) | $\Delta\tau_{\mathrm{FRET,\,syst}}$ (ns) | $\tau_r \pm \Delta\tau_r$ (ns) |
|---|---|---|---|---|---|
| sGrich | 28 | 5.0 | 29 | 0.3 | **29 ± 6** |
| d$\kappa$h | 37 | 8.0 | 46 | 0.0 | **42 ± 10** |
| dGrich | 30 | 2.7 | 32 | 0.5 | **31 ± 5** |
| sPTBP | 28 | 2.3 | 30 | 3.8 | **29 ± 6** |
| sCh | 26 | 1.6 | 30 | 0.7 | **28 ± 4** |
| sNrich | 25 | 1.7 | 27 | 0.2 | **26 ± 4** |
| s$\kappa$h | 29 | 2.0 | 31 | 0.1 | **30 ± 5** |
| dCh+ | 26 | 1.9 | 32 | 1.3 | **28 ± 5** |
| dCh− | 23 | 0.2 | – | – | **23 ± 3** |
| s$\kappa$l | 23 | 0.7 | 25 | 0.8 | **24 ± 4** |
| sNh+ | 27 | 2.1 | 32 | 4.3 | **30 ± 6** |
| dArich | 22 | 0.4 | 25 | 0.1 | **24 ± 3** |
| dErich | 25 | 1.2 | 30 | 0.4 | **28 ± 4** |
| dTRBP | 24 | 1.5 | 27 | 2.5 | **26 ± 5** |
| d$\kappa$l | 21 | 1.5 | – | – | **21 ± 3** |
| sNh− | 30 | 1.5 | 36 | 1.2 | **33 ± 5** |

Table S3: Reweighting results. Columns report the initial and reweighted means of interdye distance $r$, FRET efficiency $E$, orientation factor $\kappa^2$, and the variance of $E$. Experimental targets are listed as $E_{\mathrm{exp}}$ and $\mathrm{Var}(E)_{\mathrm{exp}}$. Additional diagnostics include $\chi^2$, the entropy change $\Delta S$, the "temperature" parameter $\Theta$, and the number of frames used $N_{\mathrm{eff}}$.

| Linker | $r_{\mathrm{init}}$ (nm) | $r_{\mathrm{rew}}$ (nm) | $E_{\mathrm{init}}$ | $E_{\mathrm{rew}}$ | $E_{\mathrm{exp}}$ | $\kappa^2_{\mathrm{init}}$ | $\kappa^2_{\mathrm{rew}}$ | $\mathrm{Var}(E)_{\mathrm{init}}$ | $\mathrm{Var}(E)_{\mathrm{rew}}$ | $\mathrm{Var}(E)_{\mathrm{exp}}$ | $\chi^2$ | $\Delta S$ | $\Theta$ | $N_{\mathrm{eff}}$ |
|---|---|---|---|---|---|---|---|---|---|---|---|---|---|---|
| sGrich | 3.55 | 3.55 | 0.90 | 0.90 | 0.87 | 0.63 | 0.63 | 0.03 | 0.03 | 0.03 | 0.81 | 0.00 | 1.17e6 | 1.00 |
| d$\kappa$h | 4.10 | 4.33 | 0.84 | 0.80 | 0.77 | 0.67 | 0.67 | 0.05 | 0.06 | 0.06 | 1.00 | 0.01 | 49.16 | 0.99 |
| dGrich | 4.07 | 4.41 | 0.82 | 0.76 | 0.73 | 0.63 | 0.64 | 0.06 | 0.08 | 0.08 | 1.00 | 0.02 | 41.91 | 0.98 |
| sPTBP | 5.09 | 5.09 | 0.67 | 0.67 | 0.66 | 0.67 | 0.67 | 0.10 | 0.10 | 0.09 | 0.27 | 0.00 | 1.03e20 | 1.00 |
| sCh | 4.24 | 5.11 | 0.82 | 0.67 | 0.64 | 0.67 | 0.68 | 0.05 | 0.09 | 0.10 | 1.13 | 0.18 | 16.52 | 0.84 |
| sNrich | 5.68 | 5.68 | 0.58 | 0.58 | 0.60 | 0.66 | 0.66 | 0.09 | 0.09 | 0.10 | 1.15 | 0.00 | 1.26e6 | 1.00 |
| s$\kappa$h | 4.36 | 5.27 | 0.79 | 0.63 | 0.60 | 0.65 | 0.67 | 0.05 | 0.09 | 0.10 | 1.75 | 0.24 | 13.94 | 0.78 |
| dCh+ | 4.57 | 5.34 | 0.75 | 0.62 | 0.59 | 0.66 | 0.67 | 0.07 | 0.10 | 0.11 | 1.35 | 0.11 | 23.41 | 0.90 |
| dCh− | 6.10 | 6.10 | 0.52 | 0.52 | 0.50 | 0.69 | 0.69 | 0.11 | 0.11 | 0.13 | 3.03 | 0.00 | 0.00 | 1.00 |
| s$\kappa$l | 5.64 | 6.06 | 0.59 | 0.52 | 0.49 | 0.64 | 0.65 | 0.11 | 0.12 | 0.13 | 1.41 | 0.02 | 54.87 | 0.98 |
| sNh+ | 5.59 | 5.95 | 0.60 | 0.54 | 0.51 | 0.66 | 0.66 | 0.11 | 0.11 | 0.13 | 1.64 | 0.02 | 64.85 | 0.98 |
| dArich | 6.05 | 6.24 | 0.52 | 0.49 | 0.47 | 0.65 | 0.65 | 0.10 | 0.11 | 0.13 | 1.30 | 0.01 | 47.88 | 0.99 |
| dErich | 5.51 | 6.05 | 0.60 | 0.51 | 0.48 | 0.65 | 0.65 | 0.11 | 0.12 | 0.13 | 1.48 | 0.04 | 40.49 | 0.97 |
| dTRBP | 6.30 | 6.41 | 0.49 | 0.47 | 0.46 | 0.66 | 0.66 | 0.11 | 0.11 | 0.13 | 1.12 | 0.00 | 49.48 | 1.00 |
| d$\kappa$l | 6.60 | 6.57 | 0.43 | 0.44 | 0.44 | 0.64 | 0.64 | 0.10 | 0.12 | 0.14 | 1.02 | 0.02 | 20.51 | 0.98 |
| sNh− | 6.42 | 6.77 | 0.47 | 0.43 | 0.42 | 0.67 | 0.67 | 0.10 | 0.12 | 0.14 | 1.11 | 0.06 | 12.86 | 0.95 |
